# Integrative genotyping and analysis of canine structural variation using long-read and short-read data

**DOI:** 10.1101/2025.03.05.641690

**Authors:** Peter Z. Schall, Jeffrey M. Kidd

**Affiliations:** Department of Human Genetics, University of Michigan, Ann Arbor, MI 48109, USA; Department of Computational Medicine and Bioinformatics, University of Michigan, Ann Arbor, MI 48109, USA

**Author notes:** **Corresponding author:** Jeffery M. Kidd, Ph.D.

**Keywords:** structural variants, long-read sequencing, mobile elements

## Abstract

Structural variation makes an important contribution to canine evolution and phenotypic differences. Although recent advances in long-read sequencing have enabled the generation of multiple canine genome assemblies, most prior analyses of structural variation have relied on short read sequencing. To offer a more complete assessment of structural variation in canines, we performed an integrative analysis of structural variants present in 12 canine samples with available long-read and short-read sequencing data along with genome assemblies. Use of long-reads permits the discovery of heterozygous variation that is absent in existing haploid assembly representations while offering a marked increase in the ability to identify insertion variants relative to short-read approaches. Examination of the size spectrum of structural variants shows that dimorphic LINE-1 and SINE variants account for over 45% of all deletions and identified 1,410 LINE-1s with intact open reading frames that show presence-absence dimorphism. Using a graph-based approach, we genotype newly discovered structural variants in an existing collection of 1,879 resequenced dogs and wolves, generating a variant catalog containing a 56.5% increase in the number of deletions and 705% increase in the number of insertions previously found in the analyzed samples. Examination of allele frequencies across admixture components present across breed clades identified 283 structural variants evolving with a signature of selection.

**Significance statement:** Existing studies of structural variation have focused on genomes sequenced with short-read technology. In this study, we systematically combine short-read and long-read data sets to identify and genotype structural variation across canines, resulting in an expanded catalog of structural variants. Analysis of allele frequencies identified structural variants in genic regions that may have been evolving under selection across breed clades.

## Introduction

Structural variation, operationally defined as insertions, deletions, duplications, or inversions of genome sequence at least 50 base pairs (bps) in length, is a critical component of genome variation (Escaramis et al., 2015). Although fewer in number than single nucleotide polymorphisms (SNPs), structural variants (SVs) typically affect a larger percentage of the genome sequence due to their larger size (Pang et al., 2010, Genomes Project et al., 2015), and have been linked as a source of genetic diversity underpinning genome evolution, environmental adaptation, and differences in gene expression (Hujoel et al., 2022, Ho et al., 2020).

The past 15 years have seen marked improvement in algorithms to identify and genotype SVs, mostly focusing on widely prevalent short-read data from the Illumina platform (Mahmoud et al., 2019). These approaches rely upon split-read and discordant read-pair signatures obtained from aligning short-read data to an existing genome reference assembly (Ho et al., 2020, Kosugi et al., 2019). A split-read occurs when a read does not have a single contiguous and complete alignment to the reference, often because the read spans the breakpoint of a structural variant. Discordant read-pairs occur when the two reads sequenced from opposite ends of a DNA fragment do not align to the reference with an orientation and apparent separation that is consistent with the bulk properties of the sequencing library. This signature indicates that the sequenced fragment may include a rearrangement breakpoint. While short-read technologies, typically paired-end reads of ∼150 bp in length, can readily identify SV breakpoints, fully resolving insertions longer than the read length remains a challenge. Identifying structural variation in genomic regions where short-reads cannot be unambiguously aligned (such as within duplications or regions with high repeat content) poses additional problems (Ho et al., 2020). With the recent advent of long-read based sequencing from Pacific Biosciences (PacBio) and Oxford Nanopore Technologies (ONT), sequenced reads can now reach lengths in the 100s of kb. This massive increase in read length has allowed for the closing of gaps in genome assemblies, enabling the identification of variants within these regions (Aganezov et al., 2022), and the full resolution of large SVs (Amarasinghe et al., 2020). Comparisons in humans and other species have shown that long-read approaches offer a more complete view of SVs, particularly of insertions longer than Illumina read lengths (Kosugi et al., 2019). Due to cost and library preparation requirements, short-read approaches continue to dominate large-scale analyses of genomic diversity. However, the development of graph-based analysis tools enables the genotyping of SVs identified from long-read data in samples with existing short-read information, offering a compelling analysis paradigm for a more comprehensive assessment of SVs (Yan et al., 2021, Chen et al., 2019).

Canines (dogs and wolves) have emerged as a powerful model for studies of genome evolution and disease. The differences in size, shape, coloration, and behavior among modern dogs makes canines an attractive system for studying the action of selection and for untangling the molecular basis of complex phenotypes (Boyko, 2011, Karlsson and Lindblad-Toh, 2008). Naturally occurring genetic variation in dogs is associated with traits, including morphological and behavior differences that have been the subject of selection (Shearin and Ostrander, 2010, Schlamp et al., 2016) as well as differences in longevity and susceptibility to multiple diseases, including cancer (Fleming et al., 2011). The canine genomics community has developed rich resources including the creation of multiple reference assemblies generated from long-read sequencing technologies (Dreger et al., 2016, Plassais et al., 2019, Freedman et al., 2016, Schall et al., 2023, Edwards et al., 2021, Sinding et al., 2021, Wang et al., 2021, Field et al., 2020, Field et al., 2022, Jagannathan et al., 2021, Player et al., 2021, Halo et al., 2021).

SVs are known to make important contributions to canine traits. Examples include deletions and duplications associated with body weight in Labrador Retrievers (Antkowiak and Szydlowski, 2023), a 133-kb duplication predisposing Ridgebacks to dermoid sinus and causing the distinctive hair ridge (Salmon Hillbertz et al., 2007), a small insertion causing globoid cell leukodystrophy in Irish Setters (McGraw and Carmichael, 2006), a *FGF4* retrogene insertion causing the short-legged phenotypes found in many breeds (Parker et al., 2009), and the well characterized copy-number variation of *AMY2B* leading to adaptation to a starch-rich diet (Axelsson et al., 2013).

Previous studies have found that dimorphic mobile element insertions make a major contribution to genomic differences among dogs (Wang and Kirkness, 2005, Halo et al., 2021). Mobile elements (also called transposable elements) are DNA sequences which can move into new genomic locations. Retrotransposons, the most abundant type of mobile element in mammalian genomes, mobilize through a “copy-and-paste” mechanism that uses an RNA intermediate that is reverse transcribed and integrated into a new genomic location (Kazazian and Moran, 2017, Levin and Moran, 2011). Retrotransposons can be further subdivided into two classes, those that contain long terminal repeats (LTRs) and those that do not. Non-LTR retrotransposons include Long INterspersed Elements (LINEs), which are autonomous retrotransposons that encode the proteins required for their mobilization, and Short INterspersed Elements (SINEs), which are non-autonomous retrotransposons that require protein(s) encoded by LINEs for mobilization. LINE-1, the autonomous retrotransposon family active across mammals, is approximately 6 kb in size and encodes two proteins, ORF1p and ORF2p, required for its own mobilization (Beck et al., 2011).

Most copies of mobile elements in the genome were inserted many millions of years ago. These sequences have acquired multiple mutations that make them incapable of mobilization and are “fixed” in the species, meaning they are found in the same genomic location in all individuals. However, some elements that were inserted more recently in evolutionary time show presence-absence variation, known as dimorphism, among individuals. These variants manifest as insertions and deletions in a sample relative to a reference genome. Several previous studies have linked mobile element insertions to canine phenotypes, including cerebellar ataxia, retinal atrophy, and merle coloration (Hitti-Malin et al., 2020, Mauri et al., 2017, Langevin et al., 2018). A recent study identified a *de novo* LINE-1 mutation as causing X-linked muscular dystrophy in a Border Collie dog, confirming that at least one type of retrotransposon remains active in the canine genome (Van Poucke et al., 2024).

In this study, we perform a comprehensive analysis of SVs in 12 canine samples with available long-read and short-read sequencing data along with genome assemblies. We compare the ability of each modality to detect SVs, highlighting the limitations of existing assemblies and of short-read data. We analyze the sequence content of identified variants, finding that dimorphic LINE-1 and SINE variants account for over 40% of all deletions. Finally, using a graph-based approach, we genotype an expanded database of SVs in an existing collection of diverse canine genomes with short-read data analyzed by the Dog10K consortium and identify SVs with frequency patterns consistent with non-neutral evolution among clades of breed dogs, including several mobile element insertions. In total, this manuscript integrates long-read and short-read genome data to provide a comprehensive view of structural variation in canines.

## Materials and Methods

### Included samples with long-read sequencing and assembled genomes

We identified a total of 12 samples that had accessible long-read sequencing data and associated assembled genomes (Supplemental Table S1). For increased readability, we refer to the different samples and assemblies by the name of the sequenced individual. The analyzed samples include 10 samples from 7 dog breeds: two Basenjis (China and Wags) (Edwards et al., 2021), two Bernese Mountain Dogs (BD and OD) (Schall et al., 2023), two German Shepherds (Mischka and Nala) (Wang et al., 2021, Field et al., 2020), a Cairn Terrier (CA611) (Schall et al., 2023), a Boxer (Tasha) (Jagannathan et al., 2021), a Great Dane (Zoey) (Halo et al., 2021), a Labrador Retriever (Yella) (Player et al., 2021, Schall et al., 2023); a Dingo (Sandy) (Field et al., 2022); and a Greenland Wolf (mCanLor) (Sinding et al., 2021). Illumina short-read sequencing was also available for each sample, except one Basenji (China). The effective long-read coverage of each sample, based on the median coverage reported by sniffles2 at identified SVs, ranges from 7x-82x (median=27.5). Effective short-read coverage was calculated based on coverage at sites included on the Illumina CanineHD BeadChip genotyping array as in (Meadows et al., 2023), and ranges from 20x-73x (median=31x). A detailed summary of the analyses undertaken in this manuscript can be found in Supplemental Figure S1.

### Canine structural variant calling

#### Long-read SV detection

Structural variants were identified from Oxford Nanopore Technologies (ONT) and PacBio long-reads using a method adapted from (Lemay et al., 2022). Reads were aligned to the UU_Cfam_GSD_1.0_ROSY genome reference with minimap2 (v2.26, RRID: SCR_018550) (Li, 2018) with standard settings, converting SAM output to BAM with SAMtools (v1.13, RRID:SCR_002105) (Li et al., 2009). Due to the incomplete nature of the assembled Y chromosome, our results are limited to assembled chromosomes (chr1-38 and X). Initial structural variants were called using sniffles2 (v2.4, RRID:SCR_017619) (Smolka et al., 2024) with a maximum deletion length of 10 Mb. To increase sensitivity, the coordinates of known tandem repeats were downloaded from the UCSC Genome Browser (RRID:SCR_005780) (Meyer et al., 2013) and included in the sniffles2 analysis. For each sample, a filtering step was applied removing: BND variants (breakend variants; where the SV breakpoints are not fully resolved), homozygous reference variants, insertions with unassembled or gap sequences (‘N’), variants less than 50bp, and variants greater than 10Mb. Subsequently, the filtered VCF files were sorted, normalized, and filtered for duplicate entries with BCFtools (v1.12, RRID:SCR_005227) (Danecek et al., 2021). Refinement of SV breakpoints and sequences was conducted with the R script breakpoint_refinement.R, which executes several additional external programs to process candidate breakpoints (Lemay et al., 2022). In short, initial local assemblies were generated for each variant with wtdbg2 (v2.5, RRID:SCR_017225) (Ruan and Li, 2020) using reads mapped +/-200bp from each SV followed by realignment of the same reads to each assembled sequence with minimap2, and the local assembly aligned to the region of the reference genome with AGE (v0.4, RRID:SCR_005253) (Abyzov and Gerstein, 2011). Since Nala and Sandy have both ONT and PacBio long-read sequencing data, an additional step of processing was applied to create a single long-read callset for each sample: the post-breakpoint refinement VCF files were merged with SURVIVOR (v1.0.7, RRID:SCR_022995) (Jeffares et al., 2017), with the following flags: 100 0 1 0 0 50. To facilitate genotyping SVs with short-read data, the final VCF files for each of the 12 samples were ported to SVmerge from the SVanalyzer package (v0.36) (https://github.com/nhansen/SVanalyzer) with the following flags: *-maxdist 15 -reldist 0*.*1 -relsizediff 0*.*1 -relshift 0*.*1 -seqspecific*. The SVmerge output VCF was processed with the R script merge_realigned_variants.R (Lemay et al., 2022), wherein SVs were selected from clusters, resulting in the final representative SV per loci.

#### Short-read SV detection

Structural variants were also identified using Illumina short-read data. Read pre-processing and alignment was conducted using the dogmap pipeline (https://github.com/jmkidd/dogmap) (Meadows et al., 2023). In summary, the pipeline consists of mapping with bwa-mem2 (v2.1, RRID: SCR_022192) (Li, 2013), followed by marking and sorting duplicates with MarkDuplicates, base score recalibration with BaseRecalibrator and ApplyBQSR, from the GATK suite of tools (v4.2, RRID:SCR_001876) (DePristo et al., 2011). Structural variants were identified from the resulting alignment files using Manta (v1.6.0, RRID:SCR_022997) (Chen et al., 2016), using standard settings. The initial Manta SV calling of Sandy failed to complete, with further investigation noting an aberrant insert-size distribution. This necessitated an additional filter to remove those reads with an apparent fragment size less than 20; this filtering ameliorated the issue. Manta SV calls were merged with svimmer (v0.1) (https://github.com/DecodeGenetics/svimmer), with standard settings, and genotyped using GraphTyper2 (v2.7.7) (Eggertsson et al., 2019). Additionally, to resolve large insertions that could not be rectified by the short-read sequencing data, structural variant genotypes were determined for the variants identified by long-reads (the output of SVmerge described above) from mapped Illumina data using Paragraph (v2.4a) (Chen et al., 2019). Within each sample, the output of the short-read manta SV calls and paragraph genotyping of long-read variants was merged with SURVIVOR, as previously described, filtering to include unique SVs only and retaining the genotype calls from Paragraph when SVs matched.

#### SVs identified through genome assembly comparisons

The third method for calling SVs consisted of comparing the published assemblies for each sample to the UU_Cfam_GSD_1.0_ROSY genome assembly. The genome alignment was conducted using minimap2 (v2.26) using the following settings: *-a -x asm5 --cs -r2k*. Resultant alignment output was sorted and indexed using SAMTools. Structural variants were called using the haploid caller from SVIM-asm (v1.0.3) (Heller and Vingron, 2021) with standard settings.

In summary, three SV call sets were generated for each sample compared to UU_Cfam_GSD_1.0_ROSY: long-read, short-read, and assembly. We additionally used short-read data to genotype SVs discovered by long-reads. To identify concordance across the three methodologies, within sample, the VCF files were merged with SURVIVOR. VCF files were then converted into table format to enable porting to R (v4.2.0, RRID:SCR_001905) (Dessau and Pipper, 2008), for processing and plotting.

### Annotation of genomic regions and structural variants

SVs were annotated using SnpEff (v5.2c, RRID:SCR_005191) (Cingolani et al., 2012), using the UU_Cfam_GSD_1.0 genomic data retrieved from Ensembl (Dyer et al., 2025), version 113, deriving genomic locations (exons, introns, intergenic, etc.) and applicable intersecting gene information. Only SVs that intersected with exons, or entire feature ablation, were considered to affect genes. Downstream analyses and annotations were limited to insertions and deletions, due to the higher confidence in calling these events compared to the other SV types.

Repetitive elements within SVs were identified using blastn (v2.10, RRID:SCR_001653) (Ladunga, 2017), comparing canine and ancestral species repeat consensus sequences retrieved from RepBase (v29.05, RRID:SCR_021169) (Bao et al., 2015). The output was then annotated to include SV length, repeat sequence length, and reciprocal percent sequence overlap between repeat sequences and SVs. Aiming to identify SVs that correspond to dimorphic repeats, we identified SVs where 1) ≥95% of the SV corresponded to a given repeat as well as 2) SVs with a 95% reciprocal overlap with a given repeat, representing near-full length dimorphic repeats. Furthermore, insertions and deletions identified as putative full-length Long INterspersed Elements (LINEs) were analyzed using BLASTX (v2.10, RRID:SCR_001653) (Ladunga, 2017) to identify intact open-reading frames for ORF1p (QOV08756.1) and ORF2p (QOV08757.1), requiring >=99% amino acid identity.

### Genotyping structural variants and selection analysis with the Dog10K dataset

The Dog10K dataset (Meadows et al., 2023) is a repository of short-read sequenced samples from nearly 2000 canines, covering well characterized breed dogs, mixed breeds, free breeding village dogs, and wolves. The output of the long-read SV pipeline, post SVmerge, was used for genotyping structural variants from the Dog10K dataset with Paragraph, with standard settings. The Dog10K dataset also included previously generated SV calls from Manta, which were retrieved and merged with the Paragraph output, as described above. Analysis of Dog10K dataset was limited to biallelic deletions and insertions on the autosomes. To ensure the inclusion of all SVs reported in the Dog10K release, genotypes of all loci were retained without regard to final Paragraph filtering status. A total of 1,879 samples were used that previously passed all variant calling filters (breed dogs=1575, mixed/other=12, village dogs=237, wolves=55). The 1,879 samples from the Dog10K project have a median coverage of 18.9x (range 12.9x-31.1x).

In the Dog10K manuscript, SNPs were used to identify signals of selection in the ancestral admixture components from 790 breed dogs that were categorized into nine broad clades (Belgian Herders, Mastiffs, Pointers, Scenthounds, Sighthounds, Spaniels, Spitz, UK Herders, and Waterdogs) using Ohana (Cheng et al., 2022). We replicated the initial generation of the allele frequency model based on the Dog10k SNP data, using the same breed-clades and number of ancestral components (K=5), and filtering parameters: minor allele frequency of 5% and no missing genotypes. We then applied the SNP-based model to the union of Manta and Paragraph SV genotypes. It should be noted that increased filtering metrics were required for the SV analysis in the Dog10K manuscript, resulting in a set of 781 dogs used for this present analysis. We applied thresholds to identify variants with allele frequency patterns inconsistent with neutral evolution as defined by Ohana. First, we applied a strict threshold of 1.67*10^−7^, corresponding to a nominal significance level of 0.05 after a Bonferroni correction accounting for the 299,115 deletion and insertions tested. Since the Bonferroni correction may be overly conservative as variant genotypes are not independent, we also applied a reduced threshold of 0.0001. A total of 283 variants passed this threshold, corresponding to the top 0.096% of all SVs evaluated.

Paragraph genotyping accuracy may be affected by variant type, size, break-point accuracy, proximity to other variants, and the presence of low-complexity or tandemly repeated sequence (Chen et al., 2019). To assess the effect of these factors on the SVs identified by Ohana, we identified insertion and deletion variants that intersected with tandem repeats in the genome reference or with alleles with greater than 70% of their sequence annotated as low-complexity or tandem repeats by SDUST or etrf (Li et al., 2020, Morgulis et al., 2006). To assess whether genotypes SVs reflect canine population structure, we performed a principal component analysis of breed dogs and wolves using smartpca (EIGENSOFT version 8.0.0)(Patterson et al., 2006). PCA was performed on one dog from each of 318 breeds as well as 55 wolves for SNPs, deletions, insertions that are not annotated as low-complexity or tandem repeats, and insertions that are annotated as low-complex or tandem repeats. PCA was limited to loci that passed all Paragraph genotyping filters. Using plink (version 1.9), loci with more than 5% missing genotypes, a minor allele frequency less than 5%, or in strong LD (−-indep-pairwise 50 10 0.1) were removed (Purcell et al., 2007).

### Generation of Figures

Unless otherwise noted, all figures were produced in R (v4.2.0), using a combination of the following R packages: ggplot2 (v3.5.1, RRID:SCR_014601) (https://ggplot2.tidyverse.org), ggpubr (v0.6.0, RRID:SCR_021139) (https://github.com/kassambara/ggpubr), ggforce (v0.4.1, RRID:SCR_022575) (https://github.com/thomasp85/ggforce), ggrepel (v0.9.5, RRID:SCR_017393) (https://github.com/slowkow/ggrepel), ggnewscale (v0.4.9) (https://doi.org/10.5281/zenodo.2543762), ComplexHeatmap (v2.20.0, RRID:SCR_017270) (Gu et al., 2016), and paletteer (v1.6.0) (https://github.com/EmilHvitfeldt/paletteer).

## Results

### Canine structural variant detection

We identified structural variants from 12 canine samples using three modalities: genome assembly, long-read sequencing, and short-read sequencing (Figure 1, Supplemental Table S1). All structural variants were identified compared to the UU_Cfam_GSD_1.0_ROSY (Mischka) genome, which is derived from a female German Shepherd Dog supplemented by partially assembled Y-chromosome sequences from a Labrador retriever. The Dog10K Consortium used the same assembly (Meadows et al., 2023). A detailed flowchart displaying the pipeline used for this manuscript can be found in Supplemental Figure S1. Regardless of input datatype, deletions and insertions were the most frequent class of detected structural variants, accounting for an average of 52.7% and 46.8% of the total calls, respectively.

**Figure 1.**
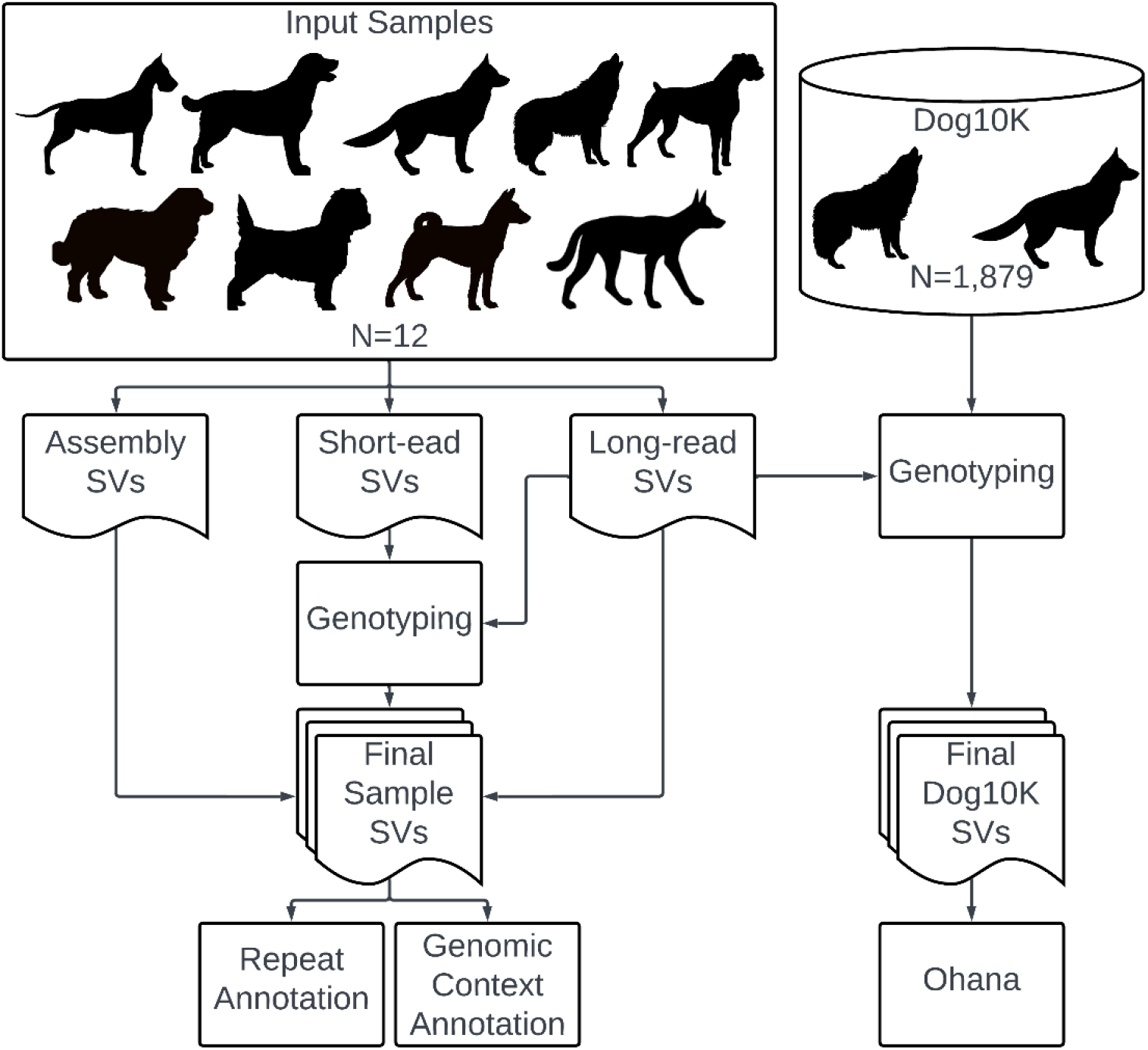
Overview of SV detection in canines. The flowchart describes integrative analysis and detection of structural variants using genome assembly, long-read, and short-read data from 12 samples. Additionally, genotyping of structural variants detected using long-reads in the Dog10K repository is depicted. All variants were identified relative to the UU_Cfam_GSD_1.0_ROSY genome assembly. Created in Lucidchart www.lucidchart.com

### Structural variation identified by assembly comparison

The number of structural variants identified by assembly comparison, which due to the self-comparison resulted in zero Mischka calls, is dominated by deletions (mean=21,509) and insertions (mean=20,712), followed by inversions (mean=68), and duplications (mean=18) (Table 1, Figure 2 Panel A, Supplemental Figure S2, Supplemental Table S2). Querying the total number of impacted base pairs by sample found an average of 21.9 Mb for deletions, 12.2 Mb for insertions, 4.6 Mb for duplications, and 17.4 Mb for inversions (Table 1, Figure 2 Panel B, Supplemental Figure S2, Supplemental Table S2). The length of SVs for both insertions and deletions displayed a similar distribution, with a mostly lefthanded dominance showing a prominent peak at ∼200-250bp and a smaller peak at ∼6,000bp (Figure 2 Panel C). The length distribution of both duplications and inversions displayed a lefthand bias (Supplemental Figure S2). SVs were filtered to include those confirmed by at least one other modality to avoid assembly errors, intersected with coding regions, and limited to genes with an annotated gene symbol. Deletions intersected with a median of 42 genes per sample (range=4-66 genes). Of the total 127 intersecting genes, 37.8% were variant in a single sample while only one, *ZCCHC10*, contained a deletion overlapping with an exon in all samples, excluding the self-comparison of Mischka. Total genes with insertions were less, numbering 70, with a median of 13 per sample (range=1-20 genes). Approximately 57.1% of the genes were variable in a single sample, and none were present in all samples (Supplemental Table S4).

**Table 1.** Summary statistics of quantified SVs and impacted base pairs in 12 canine samples by modality

|  | SV Type | Number of SVs |  |  | Basepairs (Mb) |  |  |
| --- | --- | --- | --- | --- | --- | --- | --- |
|  |  | Mean | Min | Max | Mean | Min | Max |
| Assembly | Deletions | 21,509 | 14,775 | 32,112 | 21.9 | 12.8 | 33.4 |
|  | Insertions | 20,712 | 12,686 | 28,262 | 12.2 | 8.6 | 17.7 |
|  | Inversions | 68 | 19 | 210 | 17.4 | 0.0 | 160.0 |
|  | Duplications | 18 | 7 | 34 | 4.6 | 0.0 | 44.8 |
| Long-read | Deletions | 28,148 | 16,021 | 39,021 | 12.4 | 5.3 | 16.9 |
|  | Insertions | 30,798 | 25,992 | 45,520 | 26.1 | 12.9 | 102.6 |
|  | Inversions | 205 | 86 | 289 | 13.8 | 0.5 | 147.9 |
|  | Duplications | 26 | 7 | 45 | 2.1 | 0.2 | 53.9 |
| Short-read | Deletions | 20,121 | 12,634 | 24,369 | 12.2 | 12.8 | 33.4 |
|  | Insertions | 6,019 | 732 | 9,361 | 12.2 | 8.6 | 17.7 |
|  | Inversions | 92 | 4 | 114 | 17.4 | 0.0 | 160.0 |
|  | Duplications | 232 | 110 | 319 | 4.6 | 0.0 | 44.8 |
| Short-read + Paragraph | Deletions | 25,525 | 15,542 | 36,853 | 14.0 | 6.1 | 20.7 |
|  | Insertions | 23,730 | 9,646 | 40,930 | 9.9 | 4.9 | 14.4 |
|  | Inversions | 92 | 4 | 114 | 111.0 | 9.1 | 167.9 |
|  | Duplications | 234 | 110 | 323 | 1.3 | 0.5 | 2.0 |

**Figure 2.**
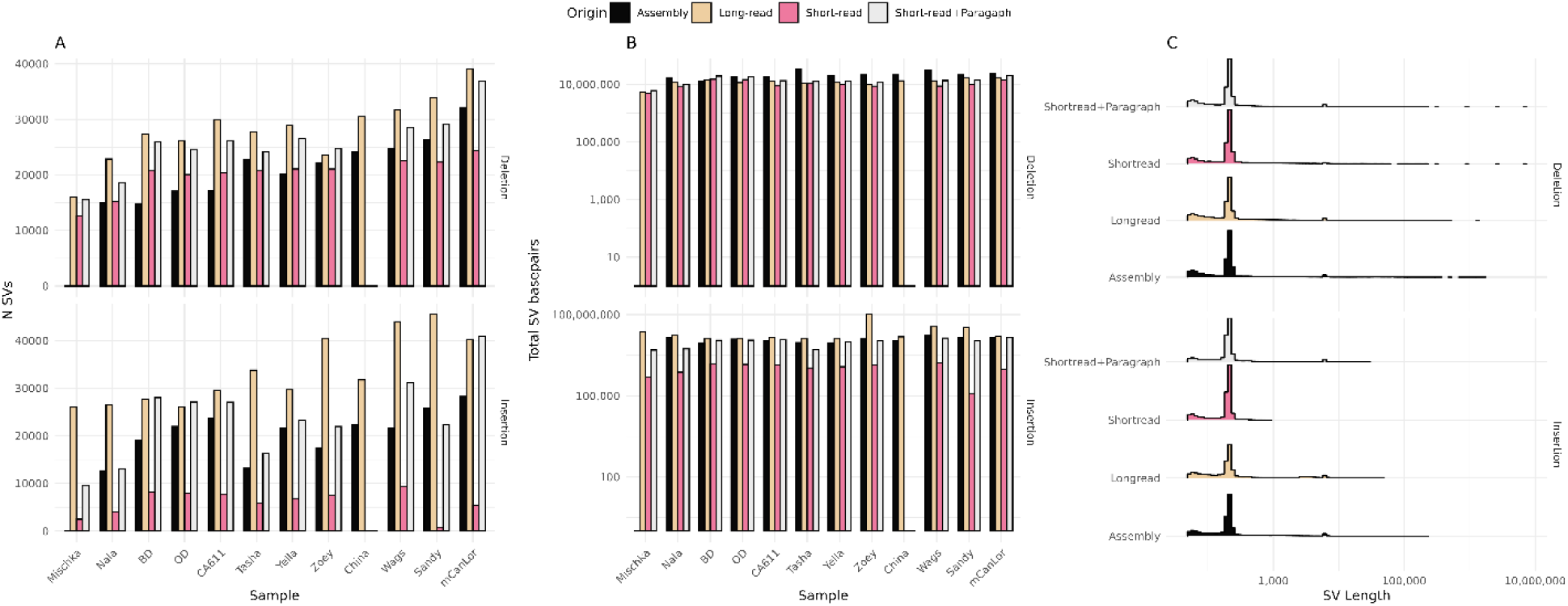
Global quantification of deletions and insertions using different data types. Panel A plots the number of deletions and insertions found across the 12 samples and four different datatypes: assembly, long-read, short-read, and short-read+paragraph. Panel B displays the summation of the total impacted basepairs. Panel C plots the distribution of SV lengths. The fill color of each bar denotes datatype origin as noted in legend.

### Structural variation identified from long-reads

Similar to assembly-based calls, the greatest number of SVs identified from long-reads were deletions (mean=28,148) and insertions (mean=30,798), trailed by inversions (mean=205) and duplications (mean=26) (Table 1, Figure 2, Supplemental Figure S2, Supplemental Table S2). Average impacted basepairs by the different types of SVs totaled 12.4 Mb for deletions, 26.1 Mb for insertions, 13.8 Mb for inversions, and 2.1 Mb for duplications. The increase in the number of SVs from long-read-based calls compared to assembly comparison can largely be attributed to heterozygous SVs. On average, 51% of deletions (range=25-99%) and 65% of insertions (range=45-98%) were heterozygous (Supplemental Figure S3, Supplemental Table S3). As expected, the greatest proportion of heterozygous calls were found in Mischka. Zoey also displayed an increased number of heterozygous calls, possibly due to long-read data quality. The length distribution of deletions, insertions, and inversions were in line with assembly calls; however, the peak of the duplications was shifted rightward.

A median of 76 genes were impacted by deletions per sample (range=15-93 genes), with a total of 241 genes affected across the samples. Approximately 42% of the impacted genes were deleted in only a single sample. Only two genes were found to have exonic deletions in all samples: *ZCCHC10* and *EEF1A1*. A median of 32 genes per sample contained exonic insertions (range=3-157 genes), affecting a total of 364 genes. The majority (84.15%) of the genes were variable in a single sample and none present in all samples (Supplemental Table S4).

### Structural variants identified using short-read sequencing

Short-read calling, which did not include China due the absence of Illumina data, was initially conducted using Manta. To better ascertain longer SVs, variants found by long-reads were genotyped using short-read data from each sample with the Paragraph software. The union of these discovered and genotyped SVs were defined as the short-read+Paragraph call set. The short-read deletions displayed a similar magnitude as previous methods (mean=20,121 per sample), with a reduced representation of insertions (mean=6,019), followed by duplications (mean=232), and inversions (mean=92) (Table 1, Figure 2, Supplemental Figure S2, Supplemental Table S2). The SV length distribution of deletions matched those resulting from assemblies and long-reads. Insertions displayed a similar distribution among shorter SVs while lacking longer, duplications were likewise similar to the assembly calls, while inversions displayed a much more flat and wider accounting of SV lengths. On average, 47% (range=32-93%) of deletions and 60% of insertions (range=4-97%) were heterozygous (Supplemental Figure S3, Supplemental Table S3). Sandy, the sample that had an aberrant insert-size distribution, was the sample with the fewest proportion of heterozygous insertion calls, and overall, the fewest insertions. We examined the 16 candidate deletions that were larger than 500 kb (Figure 2), finding that 12 are in regions annotated as segmental duplications in the UU_Cfam_GSD_1.0 (Mischka) genome, with the final 2.9 Mb of chr8 accounting for 11 of the variants (Nguyen et al., 2024a). The candidate variants were identified by a mixture of assembly, long-read, and short-read modalities, demonstrating continued difficulty in the assembly and analysis of regions that are duplicated.

While a number of statistics show relative agreement in overall SV capture by short-reads, the union of short-read+Paragraph showed increased sensitivity, specifically for longer SVs. This was most apparent for insertions, with a median percent increase of 243% identified SVs, which had a mean length of 515bp (Figure 3, Supplemental Table S2). While less of a dramatic increase in SVs, deletion calls saw a median increase of 27%, and duplications and inversions remained relatively the same. The overall impacted base-pairs increased for deletions (mean=13.95 Mb, range=6.1-13.5 Mb) and insertions (mean=9.9 Mb, range=4.9-10.9 Mb). The distribution of heterozygous calls was in-line with other sequencing modalities with deletions averaging 51% and insertions 66%. A total of 128 genes had exonic overlapping deletions and 70 genes with insertions. The median number per sample of genes with exonic deletions was 36 (range=9-73 genes) and insertions numbered 11 (range=3-28 genes). Similar to the other modalities, a substantial proportion of deletion and insertion intersecting genes were singletons, 42.3% and 64.3%, respectively (Supplemental Table S4).

**Figure 3.**
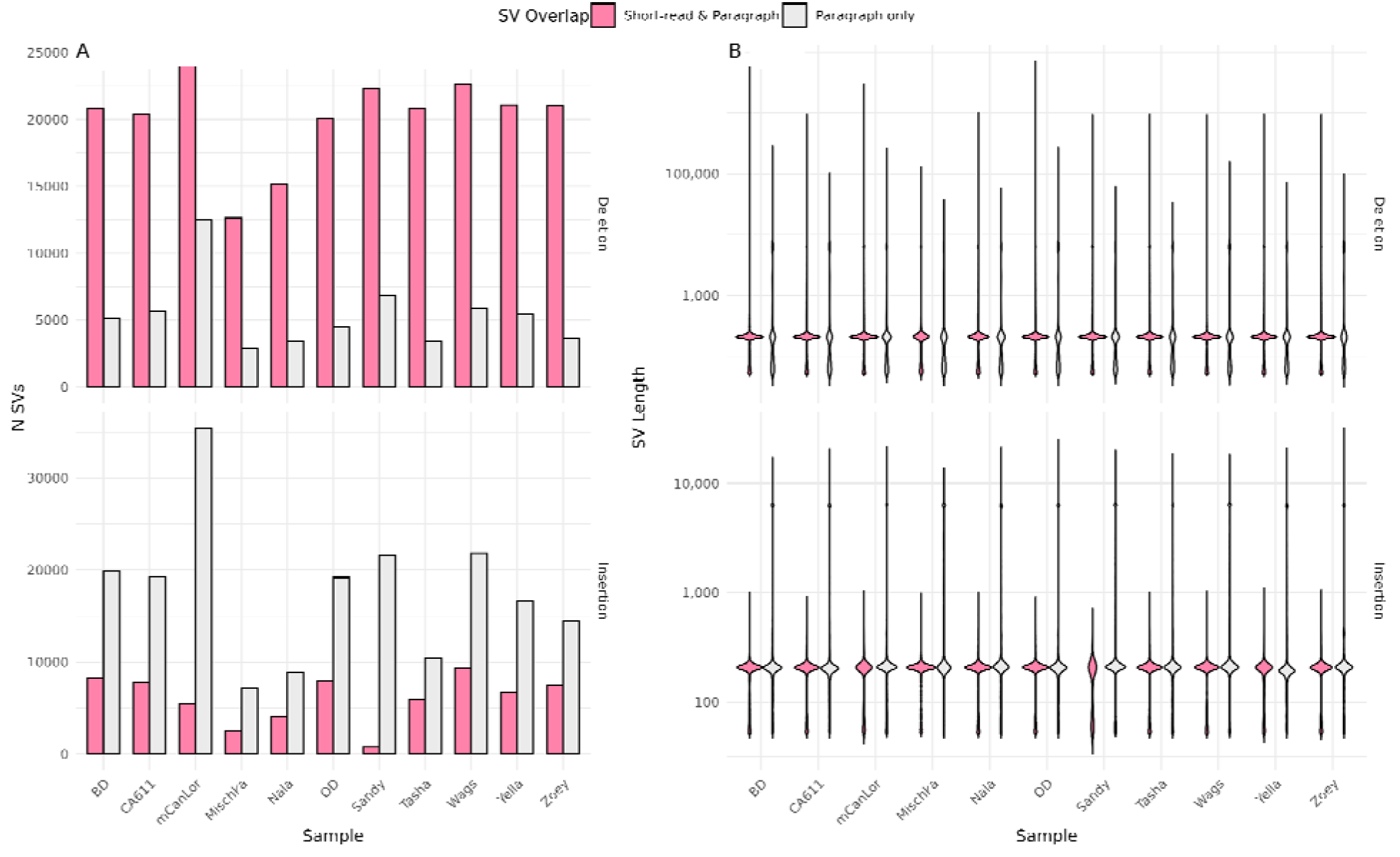
Genotyping with Paragraph enables increased detection of deletion and insertion variants using short-read data. The number and size of structural variants detected using short-read data (Manta) and those genotyped using Paragraph is shown. For both Panel A and B, the color denotes SV detection by software, pink comprised of those found jointly by the original short-read and Paragraph, while grey denotes those newly genotyped only with Paragraph. Panel A denotes the number of detected SVs, highlighting the increased number of variants that can be detected when genotyping with Paragraph. Panel B shows the length distribution of detected SVs, illustrating the ability to genotype long insertions when using Paragraph.

### Comparison of SVs by modality

Within each sample, concordance across methods was assessed by identifying the deletion and insertions variants with breakpoints within 100 bp of each other identified by each approach using SURVIVOR, e.g., long-read vs. assembly, long-read vs. short-read, long-read vs. short-read+Paragraph (Figure 4, Supplemental Table S5). We found that on average 88% of deletions and 84% of insertions discovered by comparing genome assemblies were also found in the long-read data. Conversely, on average 64% of deletions and 53% of insertions discovered in the long-read data were also found in the genome assembly comparisons. Across the samples, a total of 336,121 long-read SVs were missing in the assembly calls, of which 81.5% were heterozygous. The relatively tight grouping of both deletions and insertions found in long-read vs assembly comparison was not found when comparisons involved calls from short-read or short-read+Paragraph, with short-read modalities consistently identifying a lower fraction of insertions. However, with the inclusion of Paragraph SV calling, there is an average increase in insertion agreement compared to assembly (4%) and long-read (19%) data (Figure 4).

**Figure 4.**
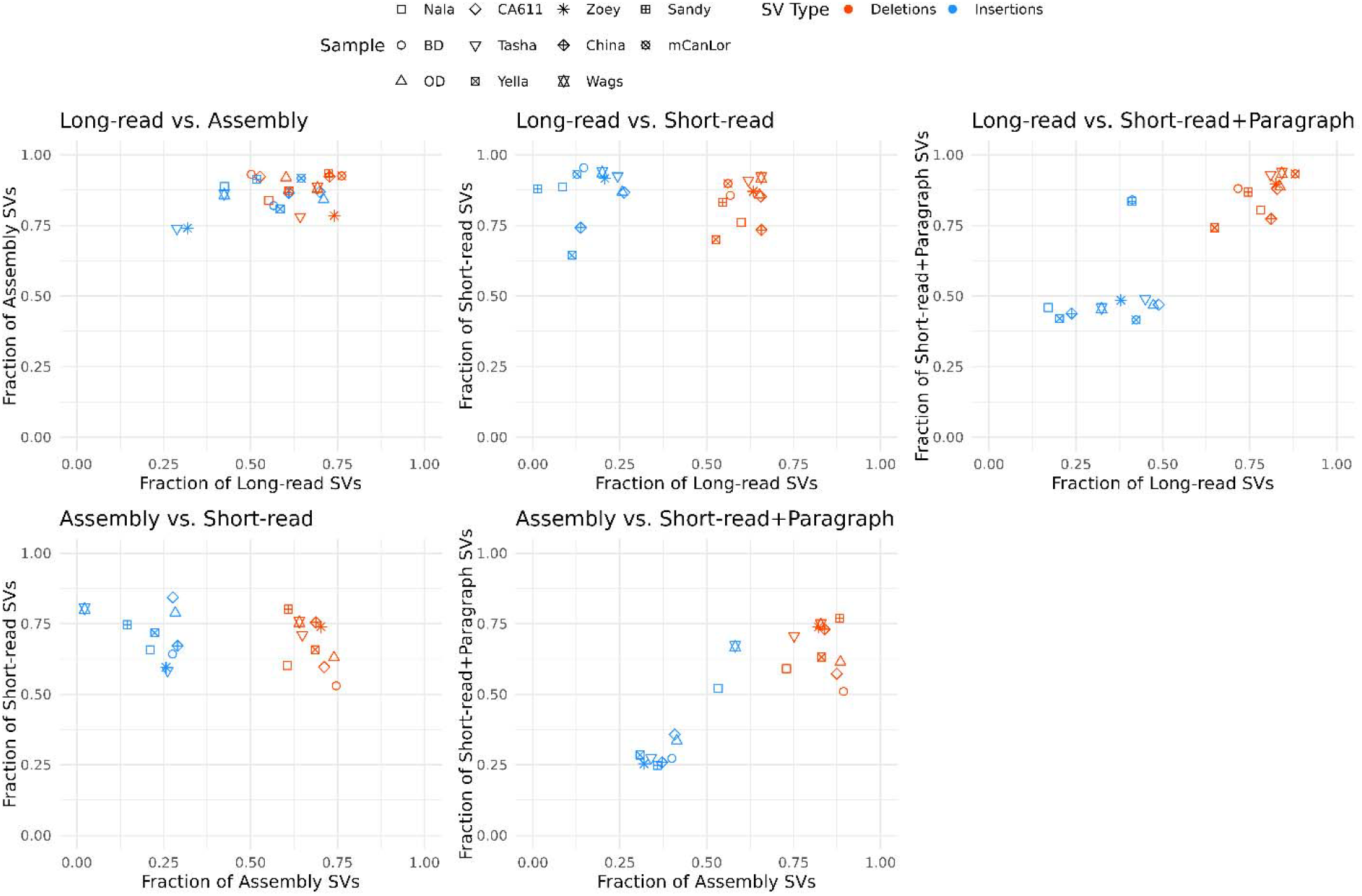
Concordance of SV detection between datatypes. Scatter plots depict the fractional quantification of detected SVs when comparing different datatypes (e.g., long-read vs. assembly, long-read vs. short-read, etc.). Each panel depicts a specific comparison with the axis quantifying the fraction shared. The shape of each point denotes the sample and color denotes SV type: deletions in red and insertions in blue.

### Repeat annotation of SVs

All identified deletion and insertion sequences, representing 148,521 and 283,895 merged deletion and insertion loci, were extracted and annotated for matches to canine and ancestral repeat sequences from RepBase using blastn. The SV-repeat data was then filtered to identify those variants with at least 95% of length annotated as a given repeat. These SVs correspond to putatively variable repeats, as opposed to larger SVs that incidentally contain repetitive sequence. This filtered dataset contained a total of 18,082 deletions and 69,103 insertions, with a mean per sample of 5,492 deletions and 6,690 insertions annotated as variable repeats (Figure 5). As expected, the repeat annotation was dominated by SINEC2A1 sequences, accounting for 72.5% of deletions and 67.4% of insertions that are annotated as repeats and 33.8% and 10.7% of the total deletions and insertions, respectively. The second largest category consisted of Long INterspersed Nuclear Elements-1 (LINE-1s), comprising 26.6% of deletions and 31.8% of insertions of the repeat annotated SVs, and 12.0% of total deletions and 5% of total insertions (Figure 5). Thus, variable SINEC and LINE-1 sequences account for 45.8% and 15.7% of all deletion and insertion structural variants detected in this dataset.

**Figure 5.**
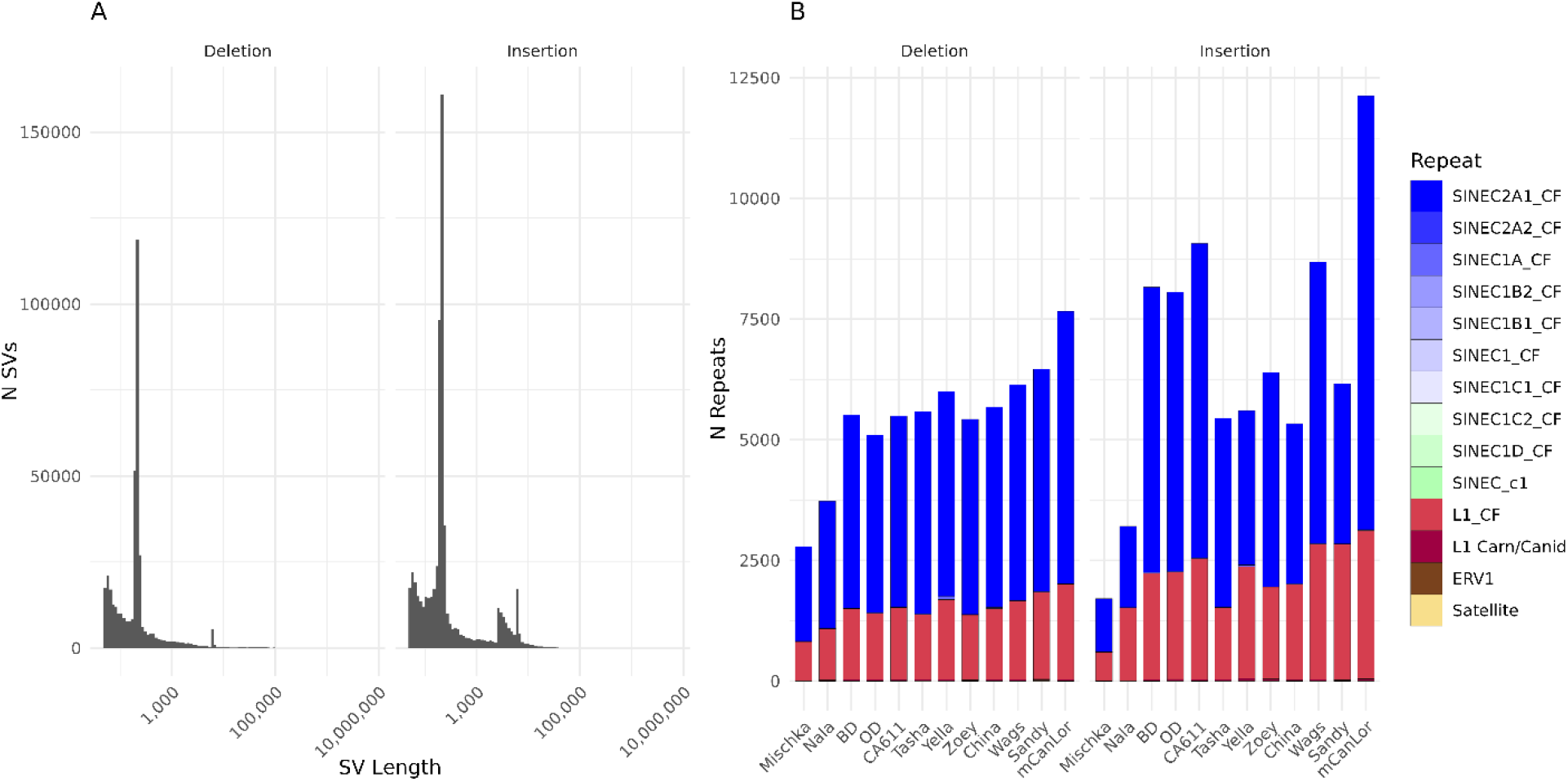
Detection of SINEs and LINEs from SV data. The length distribution of all deletions and insertions, regardless of origin datatype, are depicted in Panel A. These deletions and insertions were scanned for the presence of repeat sequences via blastn using canine and ancestral SINE and LINE sequences from RepBase. To identify those SVs comprised of a singular repeat, SVs were only included with at least 95% of the length constituting a repeat sequence. Panel B quantifies the number of detected SVs by repeat sequence name and/or family, split by either deletion or insertion.

The SV-repeat data was further filtered to identify variants putatively corresponding to full-length LINE-1s by requiring at least 95% reciprocal overlap with the element consensus sequence, which were primarily comprised of L1_CF sequences. A mean of 332 deleted and 709 inserted full-length LINE-1s were found across samples. Full-length LINE-1 sequences were further queried to identify those with intact open reading frames for the ORF1p and ORF2p proteins (>=99% sequence identity), splitting by SV type (Figure 6). A total of 1,410 variable LINEs with intact ORFs were identified with a sample median of 205 LINE-1s (Supplemental Table S6). The majority of the intact LINE-1s (82.1%) were present in a single sample, with 41.8% of the singletons originating from mCanLor, the Greenland Wolf. In fact, mCanLor had the largest number of intact LINE-1 sequences derived from SV events (n=707, deletions=233, insertions=474). Excluding Mischka, the samples with the fewest LINE-1 insertions (BD, OD, CA611, Yella, and China), were sequenced on the ONT platform only, possibly due to sequencing errors that disrupted apparent ORFs.

**Figure 6.**
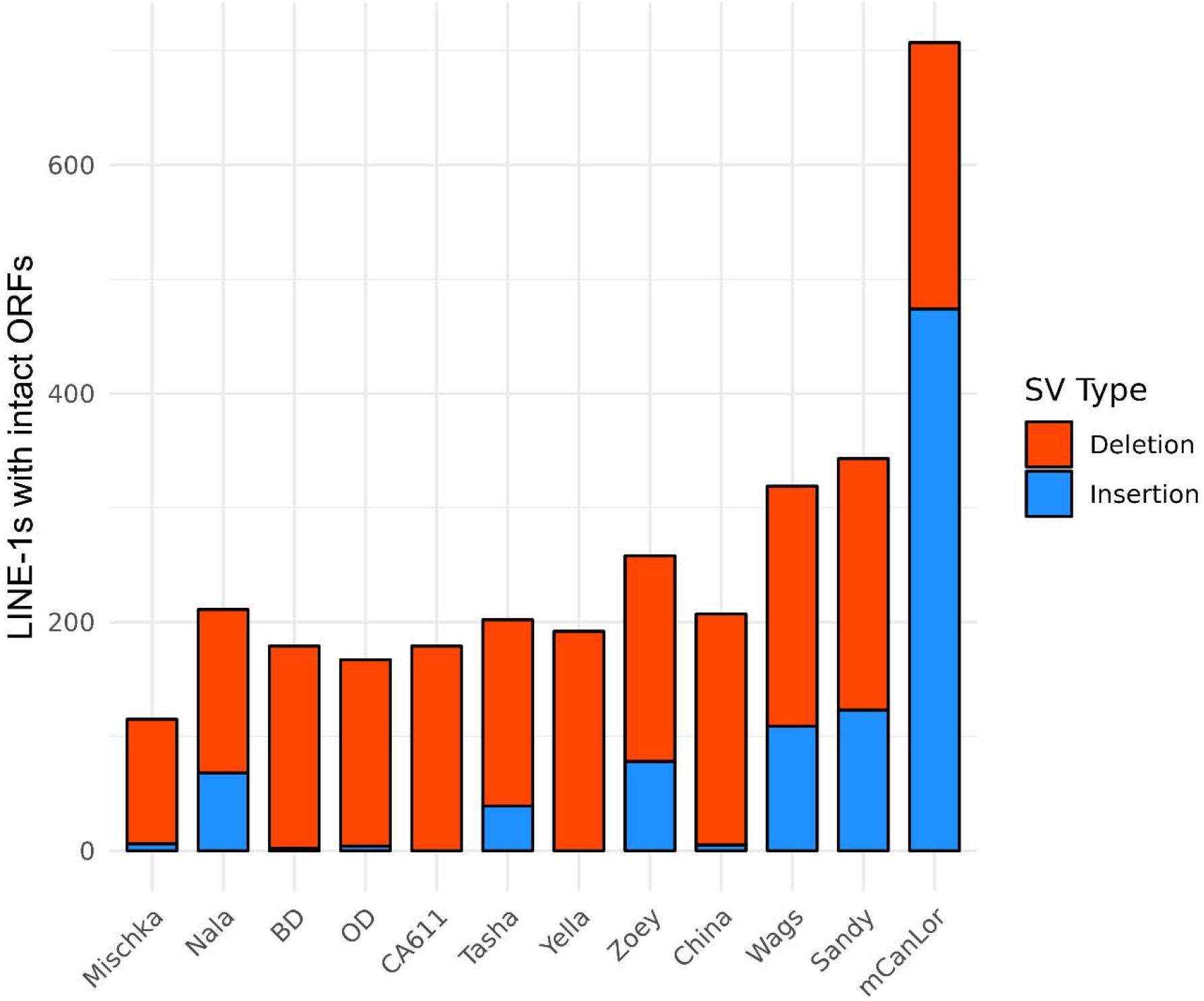
Census of dimorphic full-length LINE-1s with intact open reading frames. SVs corresponding to full-length LINE-1 sequences were extracted and scanned for intact open reading frames (>=99% sequence similarity) and quantified by either deletion or insertion for each sample. The stacked barplot lists deleted LINE-1s in red, and inserted LINE-1s in blue.

### Signals of selection in structural variant allele frequencies

We leveraged the Dog10K dataset (Meadows et al., 2023), which consists of short-read sequencing of 1,879 canines, comprised of characterized breed dogs, free breeding village dogs, and wolves to assess frequency patterns of structural variation. We applied the Paragraph method to genotype the union of the variants originally identified in the Dog10K collection and the variants identified in the 12 analyzed genomes, limiting analysis to biallelic insertions and deletions. To ensure inclusion of all SVs reported in the Dog10K release, we considered all genotyped loci without regard to the filtering status reported by Paragraph on the Dog10K data. This approach thus offers genotype information in 1,879 samples for variants discovered using both short-read and long-read data. Similar to the increase seen previously when applying the Paragraph method, we found a 56.5% increase in the number of deletions and 705% increase in the number of insertions relative to the Manta-only short-read calls (Figure 7).

**Figure 7.**
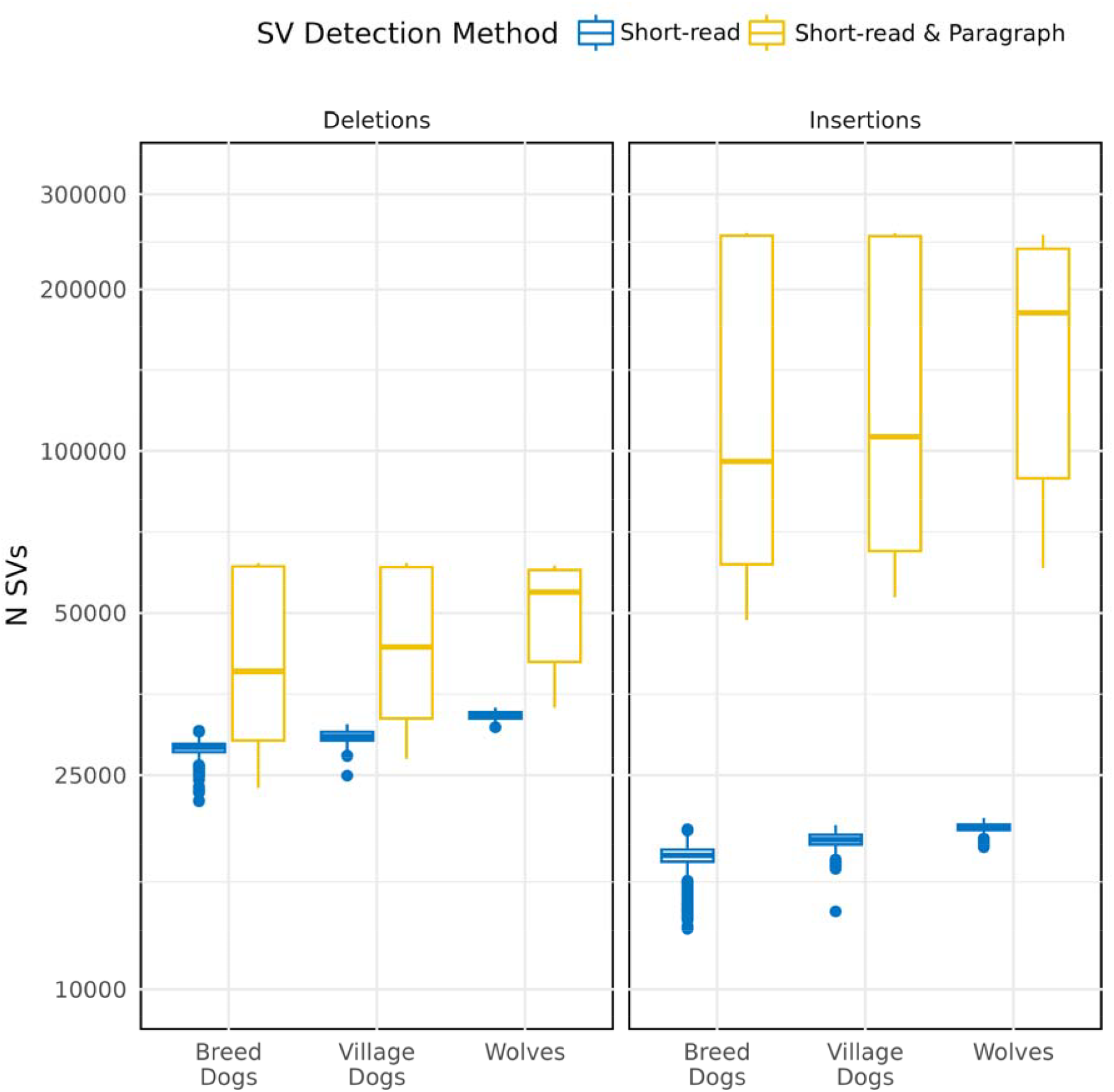
Genotyping SVs identified using longreads in the Dog10K sample collection. Box plots depict the number of deletions and insertions genotyped across the Dog10K collection using the shot-read and short-read+Paragraph approaches. The data are plotted by canine category (Breed Dogs, Village Dogs, and Wolves), as noted along the x-axis. The tabulation of the original short-read SVs is in blue, while short-read+Paragraph is in yellow.

We next used the comprehensive catalog of insertion and deletion variants genotyped in 1,879 canines to search for allele frequency profiles indicating SVs that may be evolving under selection. Previously, the Dog10K consortium identified single nucleotide variants that may have been evolving under selection among major breed clades (Meadows et al., 2023). This analysis was performed using Ohana, a program which models allele frequencies among ancestry components to identify variants with frequency profiles that are not consistent with genome-wide profiles (Cheng et al., 2022).

Following the procedure used by the Dog10K consortium, we analyzed 781 breed dogs across nine clades (Belgian Herders, Mastiffs, Scenthounds, Sighthounds, Spaniels, Spits, Pointers, UK Herders, and Waterdogs). Using SNP genotype data, we replicated the ancestry proportions identified previously and found five ancestral components that largely corresponded to specific breed clades: 0 = Collie & Shetland Sheepdog (within the UK Herding clade), 1 = Spitz, 2 = Scenthounds, 3 = Mastiffs, and 4 = Pointers & Spaniels (Supplemental Figure S4 and Figure 8). We then fit the Ohana model to biallelic SV genotype data, from a total of 299,115 deletions and insertions, to identify SVs with allele frequencies that did not fit genome wide profiles identified using SNPs in each ancestral component. A total of 81 sites that exceeded a Bonferroni-corrected significance threshold (1.67*10^−7^) were identified across the 5 ancestral components, although we note that this threshold is conservative since genotypes are not independent. The largest number of SVs were found in the ancestry component maximized in Collie & Shetland Sheepdog (n=64), followed by Mastiffs (n=10), Spitz (n=5), and Scenthounds (n=2) (Supplemental Table S7). For comparison, selection analysis was also repeated using SNP genotypes.

**Figure 8.**
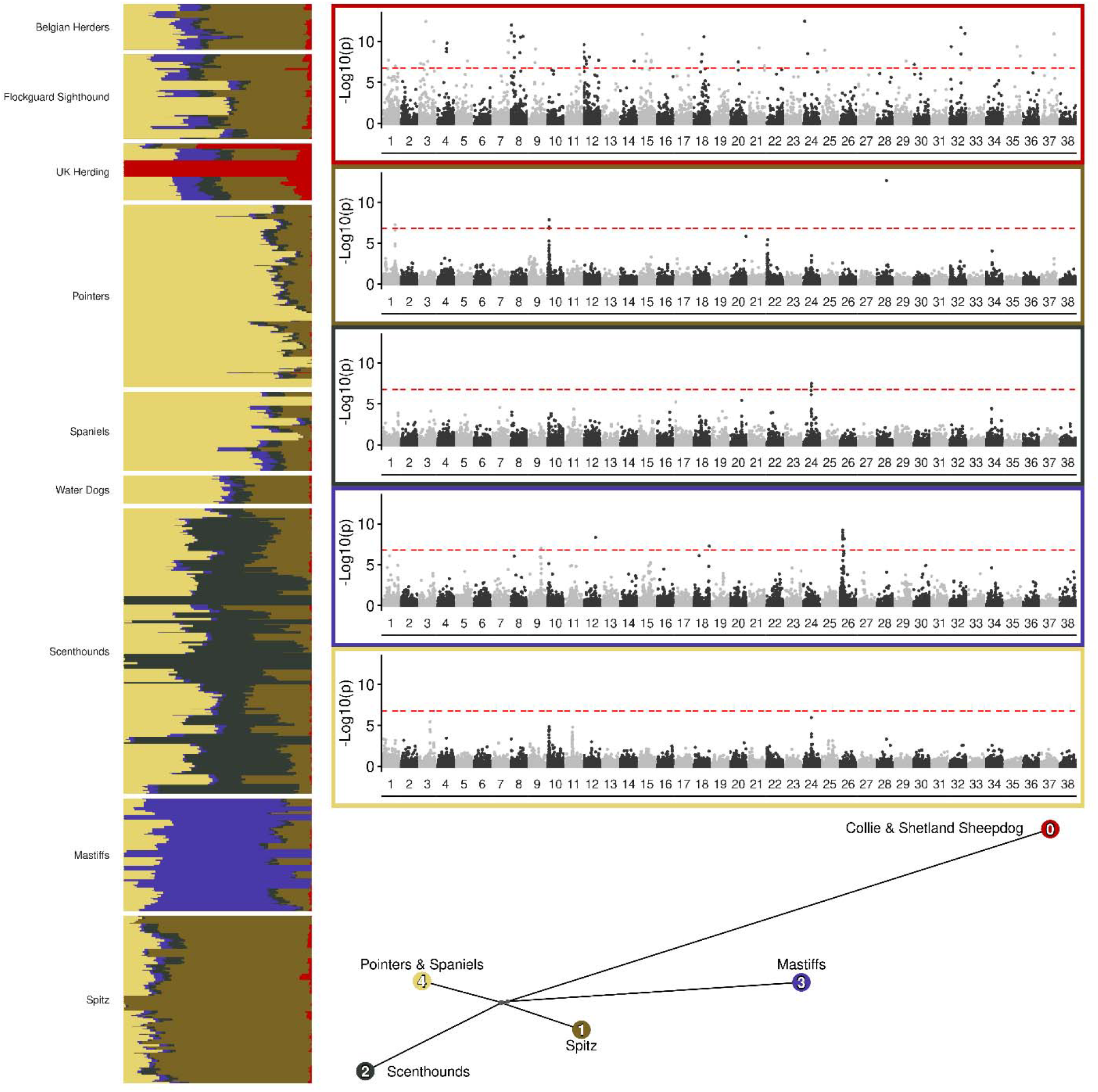
Structural variants with signatures of selection detected by Ohana with five ancestral components. Left vertical panel plots the population structure for the nine dog-breed/clade group with *K*=5 ancestral components based on the SNP data. Manhattan plots depict selection scans for SVs in each of the five ancestral components, exterior border color corresponds to color of each ancestral component, horizontal dashed red-line denotes Bonferroni threshold. The bottom network tree depicts the inferred relationships among the ancestral components from the SNP data, labeled with the breeds in which each component is maximized, filled color corresponding to both the Manhattan plot border and population structure.

Specific breeds that were maximized within the selected clades included, Spitz: Japanese Spitz, Volpino Italiano, and the Pomeranian; Scenthounds: Billy, English Foxhound, Otterhound, Great Anglo-French White and Orange Hound, and Great Anglo-French Tricolour Hound; Mastiffs: Bull Terriers, English Bulldogs, Miniature Bull Terrier, and Boxer; Pointers & Spaniels: Sussex Spaniel, Ariège Pointer, Bourbonnais Pointing Dog, English Pointer, and Braque Français.

Several of the identified SVs overlap genes previously linked to breed specific traits or disorders. Of the 64 Collie & Shetland Sheepdog SVs, 34 were located within the introns of annotated genes. This included a 7.8 kb intronic deletion within *NHEJ1*; this deletion has been previously associated with a disorder of ocular development found in Collies and related breeds (Collie eye anomaly [*cea*]) (Parker et al., 2007) (Figure 9, Supplemental Figure 5, Supplemental Table S8). We note that the identified deletion has a more significant selection signal than other markers, including SNPs, in this region.

**Figure 9.**
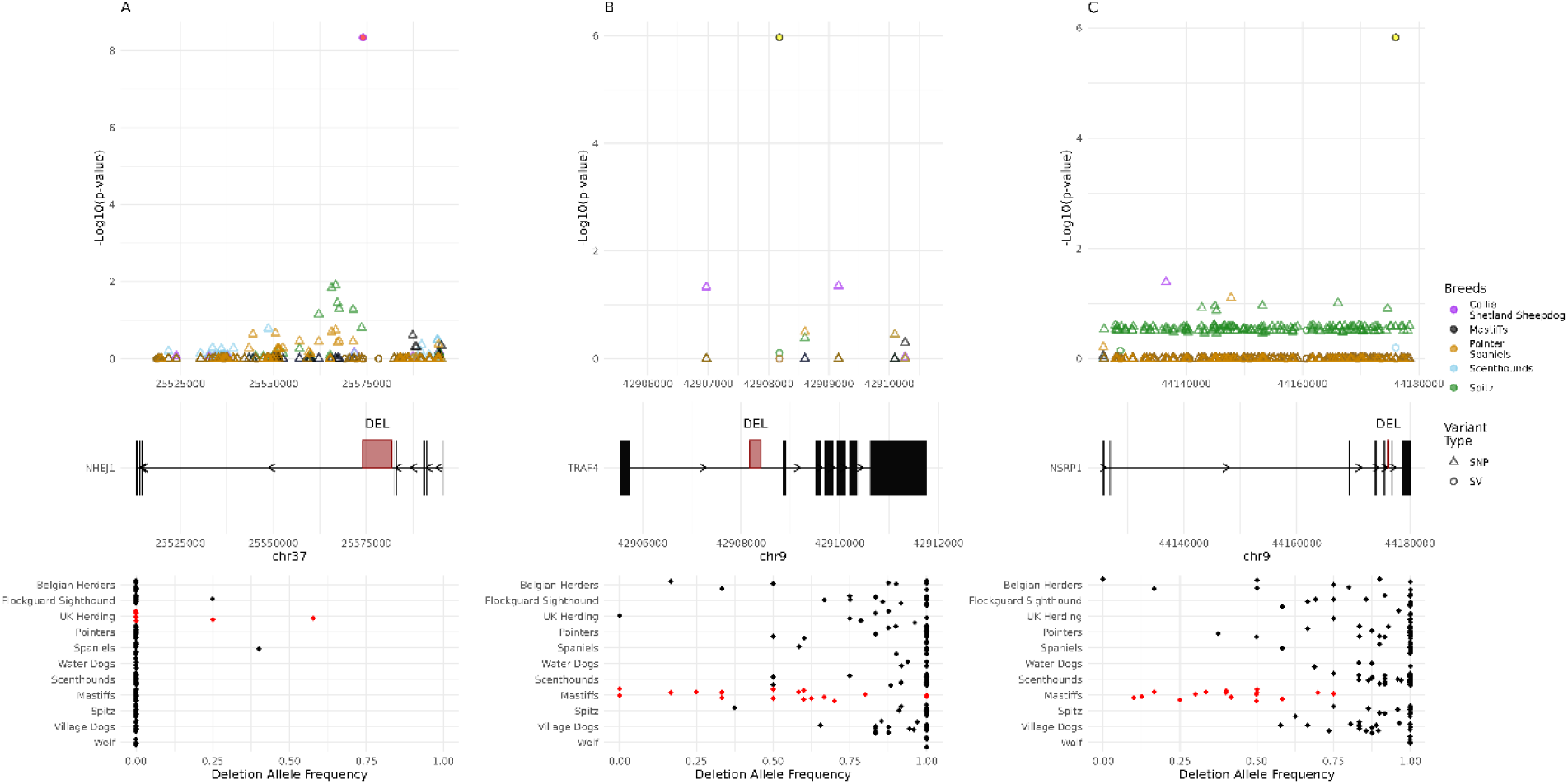
Structural variants with signals of selection. The output of Ohana, combining SNP and SV data, for three examples in their specific genomic context across the included canine breed/clade categories is depicted. Each panel consists of three sub panels: the top displaying the genomic coordinates along the x-axis and the significance along the y-axis reported by Ohana, the middle displays the relevant gene model within the region, with thick boxes denoting the position of exons and the position of most significant SV given in red, the bottom displays the allele frequency by breed or geographic location for the most significant SV. For each top panel, the color denotes the canine grouping and shape delineating variant type. Within the bottom allele frequency panel, the red colored dots correspond to the clade identified via Ohana for the associated SV selection signal. Ohana output can be found in Supplemental Table S7, and allele frequencies in Supplemental Table S8.

In the Spitz group, we found three SVs within introns of *HMGA2*, a gene which has been associated with body weight (Rimbault et al., 2013), and ear type (Webster et al., 2015), although SNPs in the region also show statistical evidence for selection (Supplemental Figure S6). SVs identified in the Mastiffs clade intersected with the introns of three genes: *ABCB9, PITPNM2*, and *DENR*, that last of which was found to have two intronic deletions and has been implicated in canine glioma susceptibility (Miller et al., 2019, Truve et al., 2016) (Supplemental Figure S7). Each of the deleted sequences were identified as SINEC2A1 repeats with intact poly(A) tails and target site duplications. All four of the SINEC sequences are present in the UU_Cfam_GSD_1.0 reference genome, derived from a German Shepherd. Consistent with the recent origin of the SINE insertions, a high deletion frequency is found across canine populations, the SINEs were absent in all available wolf samples, while found among Mastiffs. Interestingly, the two polymorphic SINEs in *DENR* have approximately 96% sequence identity and are present in an opposite orientation in adjacent introns, with one located near a coding exon, suggesting that they may affect splicing (Supplemental Figure S8).

Relaxing the level of significance to p<=0.0001, increased the total number of identified SVs to 283. This increased list included a Collie and Shetland Sheepdog intronic SV within *AP3B1*, where a homozygous loss-of-function mutations have been associated with gray Collie syndrome (Benson et al., 2003), as well as a large deletion covering multiple exons from *MX2. MX2* contributes to the innate immune defense against viruses (Haller et al., 2018, Betancor, 2023) and regulates cell-cycle progression in melanoma (Juraleviciute et al., 2020). Variation in *MX2* has also been linked to wool and skin traits in sheep (Bolormaa et al., 2021) and increased expression of *MX2* has been found in canine atopic dermatitis (Plager et al., 2012). Similarly, three intronic deletions were found below the relaxed threshold in the Spitz group within *RCL1*, a gene implicated in snout ratio and curly tail phenotype (Vaysse et al., 2011). Two additional Mastiff intronic SV deletions with signals of selection below the relaxed threshold were found within *TRAF4* and *NSRP1*. Similar to the previous three genes identified with the Mastiffs clade, each of the deleted sequences were identified as SINEC2A1 repeats with intact poly(A) tails and target site duplications and not present within the mCanLor wolf reference genome (Supplemental Figure S9). *TRAF4* plays an important role in bone development (Regnier et al., 2002) and promotes osteogenic differentiation (Li et al., 2019). Overexpression of *TRAF4* has been implicated in the regulation of tumor formation in osteosarcoma (Yao et al., 2014b, Yao et al., 2014a), a type of bone cancer that is common in certain dog breeds, including Mastiffs (Edmunds et al., 2021). *NSRP1* is a splicing regulatory protein (Kim et al., 2011). Homozygous loss-of-function alleles of *NSRP1* lead to a severe neurodevelopmental disorder (Calame et al., 2021), while *NSRP1* function has been linked to adipogenesis and modulation of weight in pigs (Liu et al., 2024, Yang et al., 2019). Mastiffs are a collection of large and giant dog breeds, with a high incidence rate of cancer and short lifespan (Bell and Hesketh, 2021).

The accuracy of SV genotypes determined by Paragraph may vary based on variant type, variant density, and sequence content, with clustered loci and low-complexity or tandem repeat loci being particularly problematic (Chen et al., 2019). Analysis of sequence content showed that 26.5% of insertions that pass all Paragraph filters are low complexity or tandem repeat sequence, while 8.8% of deletions meet the same criteria. Furthermore, 34.9% of insertion loci were located within 25 bp of another insertion, while only 10% of deletions were within the same distance of each other. A principal component analysis shows that each SV type differentiates dogs from wolves and, in an analysis limited to breed dogs, stratifies Asian breeds along PC1. PCA of deletions yields a clearer clustering of samples than observed with insertions, but separates breeds closely related to German Shepherd Dogs from the others (Supplemental Figure S10).

## Discussion

Advances in long-read sequencing have ushered in a new era of genomics and have enabled the generation of gap-free telomere-to-telomere assemblies (Marx, 2023). These resources offer a refined view of complex genome variation, including structural variation involving duplicated and repetitive sequences. Despite the power of long-read approaches, most existing catalogs of genetic variation are derived from short-read data. The canine research community has generated a rich collection of genomic resources including thousands of diverse samples sequenced using Illumina short-read sequencing along with multiple samples with long-read data. In this study, we present a uniform catalogue of SVs using a wealth of publicly available data covering short-read, long-read, and assembled genomes from 12 canine samples. Further, we used recently developed graph-based algorithms to genotype the discovered variants in a collection of 1,879 dog and wolf samples sequenced by the Dog10K consortium.

As expected, the spectrum of SVs found between genome assemblies is dominated by insertion and deletions. However, the analyzed canine assemblies are haploid representations of diploid genomes and offer an incomplete view of variation present in each sample. This is reinforced by comparison with SVs identified through alignment of the raw long-read data used to generate each assembly, where the increased number of identified SVs reflects heterozygous variants that are not represented in the respective assemblies. For most samples, an excess of heterozygous insertions was found when analyzing the long-reads, suggesting that existing assemblies may show a bias towards representing the shorter, deletion allele at heterozygous sites. Differences in data and assembly quality among the samples likely contributes to these patterns. The assemblies were generated over a period of multiple years using different sequencing technologies and assembly algorithms. Our comparisons highlight the value of approaches that comprehensively represent the variation present within diploid genomes.

The size distribution of identified insertion and deletions displayed peaks that correspond to the size of dimorphic SINE and LINE-1 elements. Comparison with repeat element consensus sequences showed that variable SINE and LINE-1 sequences from the most recent active families account for 45.8% and 15.7% of all deletion and insertion SVs. This includes 1,410 dimorphic LINE-1s that appear to have intact open reading frames for both the ORF1p and ORF2p proteins required for retrotransposition. The greatest contributor of intact LINE-1 sequences was found in the mCanLor Greenland Wolf genome, the only analyzed sample sequenced using PacBio HiFi reads, highlighting the importance of sequencing technology to the accurate base-pair level resolution of variation among repetitive sequences.

As expected, comparisons across sequencing technologies confirmed the dramatically reduced power of short-read approaches to capture insertion variants. However, variants discovered with long-reads can be genotyped using short-read data. To offer a more complete assessment of canine variation, we therefore genotyped the expanded SV catalog in samples from the Dog10K collection, resulting in a 56.5% increase in the number of deletions and 705% increase in the number of insertions previously reported in these samples (Meadows et al., 2023). The genotyping approach we employed used sequence graphs to analyze each variant locus independently. We anticipate that using a truly pangenome graph approach to assessing structural variation will become possible as high-quality, diploid genome assemblies are released from multiple canines. A pangenome approach may also yield more robust genotypes for some classes of SVs. For example, the Paragraph genotyper has reduced accuracy for tandem repeats and low-complexity sequence, a type of variation that is common in canine genomes (Laidlaw et al., 2007). Paragraph analyzes each locus independently and is known to have reduced performance when multiple variants are clustered nearby. The call set analyzed in the Dog10K samples includes many clustered insertions, with 34.9% of insertions located within 25 bp of another insertion variant. Regions with clustered or nested alleles may be better represented using a pangenome graph. The position of samples in a PCA suggests that insertion genotypes may be of lower quality than deletion genotypes. Although deletion genotypes largely recapitulate known profiles of dog population structure, breeds related to German Shepherds cluster separately. This may reflect a bias for identifying deletions relative to the UU_Cfam_GSD_1.0/canFam4 reference, which was derived from a German Shepherd Dog. In a prior analysis of SNPs, German Shepherd Dogs pull out along higher principal components, an observation that does not appear to be caused by SNP reference bias (Nguyen et al., 2024b).

The Dog10K consortium previously analyzed single nucleotide variants to identify signatures of selection in the ancestral components present across major clades of breed dogs. We repeated this analysis using the expanded catalog of 299,115 deletions and insertions, revealing examples where a genotyped insertion or deletion has a clearer signal of selection than flanking SNPs (Figure 9). To make full use of the Dog10K SV call set, we included all deletions and insertions regardless of filtration status in the Paragraph analysis. However, we note that 80/81 loci with a p-value smaller than the strict cutoff, and 279/283 of all loci with a p-vale smaller than the relaxed threshold, passed all Paragraph filters (Table S7).

Consistent with their major contribution to canine SVs, we found that 90 of the 283 variants that exceed the reduced selection threshold (p<=0.0001) corresponded to SINE or LINE deletions. Mobile element insertions can disrupt gene function through multiple mechanisms, including disruption of protein-coding exons, alterations to gene expression, and disrupted splicing processes (Hancks and Kazazian, 2016, Cordaux and Batzer, 2009). Recent reports have highlighted the potential of pairs of SINEs located in inverted orientation to disrupt splicing due to the formation of secondary structures in pre-mRNA, a mechanism implicated as contributing to the tail loss in apes (Xia et al., 2024) and the evolution of skin pigmentation variation in humans (Kamitaki et al., 2024). We identified a pair of dimorphic SINE sequences in introns of *DENR* that appear to have evolved under selection in Mastiffs and that have such an inverted orientation (Supplemental Figure S8). Additional studies will be required to determine whether improper splicing is found in dogs that carry the identified SINE insertions. Several of the genes containing SVs with signals of selection were also identified by an analysis of SNPs (Meadows et al., 2023), making it challenging to identify specific mutations that are functionally important. Several of the identified genes have been linked to diseases, making it unclear how to interpret statistical signals suggesting they were selected across breed clades. One possibility is that the identified variants are deleterious mutations that are tightly linked to variants that contribute to breed-defining phenotypes that have yet to be identified. Another possibility is that the variants have pleiotropic effects that both promote characteristics desired during breed formation while also contributing to the development of disease. More focused studies that link models of gene function with evolutionary dynamics across time periods will be required to identify the precise molecular changes underlying the formation of modern breeds.

## Data availability

VCF files describing the identified structural variants have been deposited in the Zenodo data archive under accession DOI 10.5281/zenodo.14968873. A UCSC browser track hub depicting the detected SVs available at https://github.com/KiddLab/dog-long-read-sv

