## Supplementary Figures for "Integrative genotyping and analysis of canine structural variation using long-read and short-read data"

**Supplemental Figures**

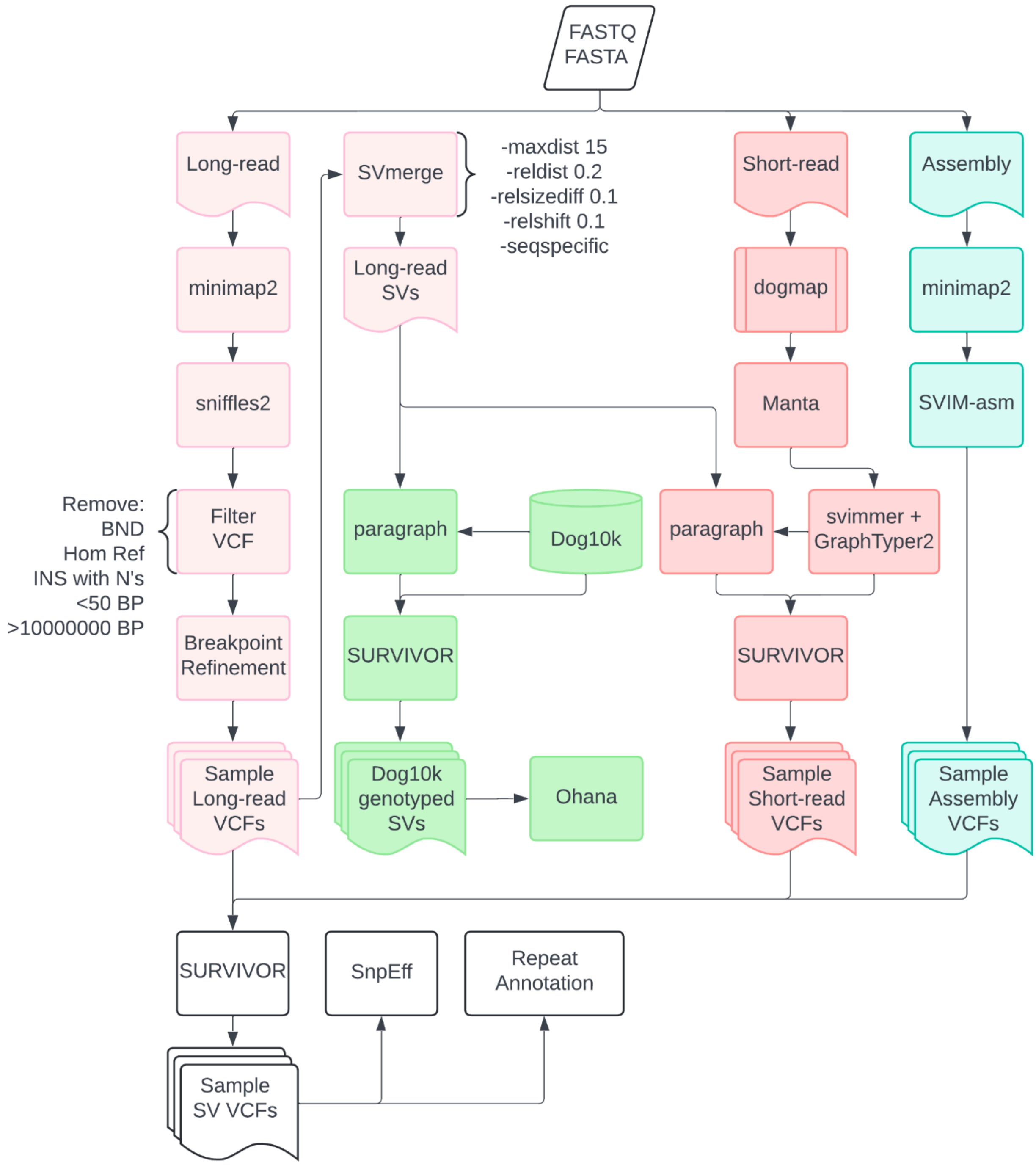


**Supplemental Figure S1 – SV detection and analysis pipeline**

The flow of analysis for the detection of structural variants for three different datatypes: long-read, assembly, and short-read. All SVs were generated compared to the UU_Cfam_GSD_1.0 genome assembly. The processing of long-read data is highlighted in light pink, short-read in dark pink, assemblies in light teal, and genotyping of Dog10K data via paragraph in green. Created in Lucidchart www.lucidchart.com.

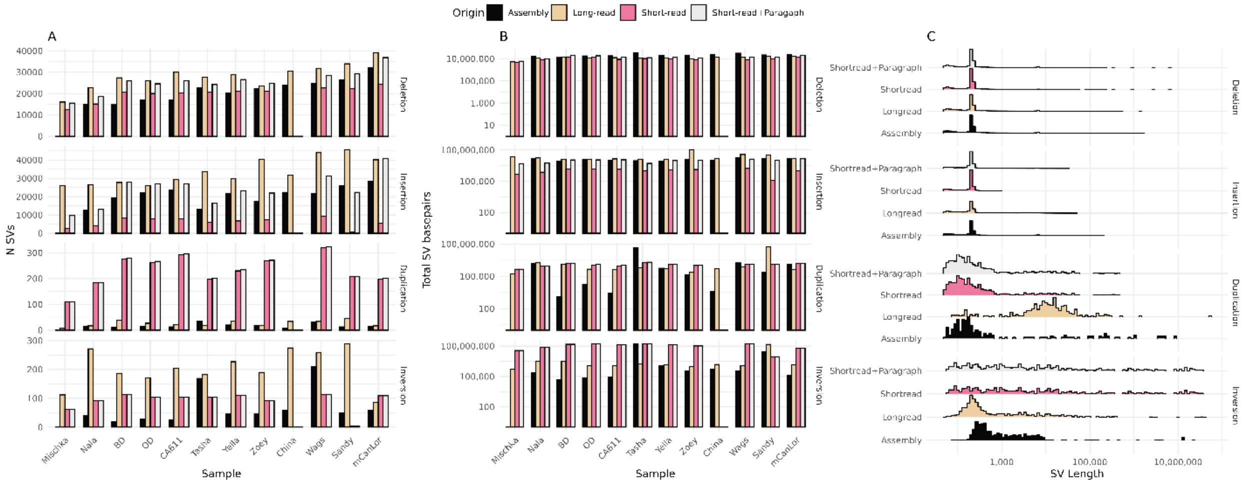


**Supplemental Figure S2 - Global quantification of SVs from samples with long-read sequencing**

Panel A plots the number of SVs found across the 12 samples and four different datatypes: assembly, long-read, short-read, and short-read+Paragraph. Panel B displays the summation of the total impacted basepairs. Panel C plots the distribution of SV lengths. The fill color of each bar denotes datatype origin as noted in legend. The top two rows detailing deletions and insertions are replicated in Figure 2.

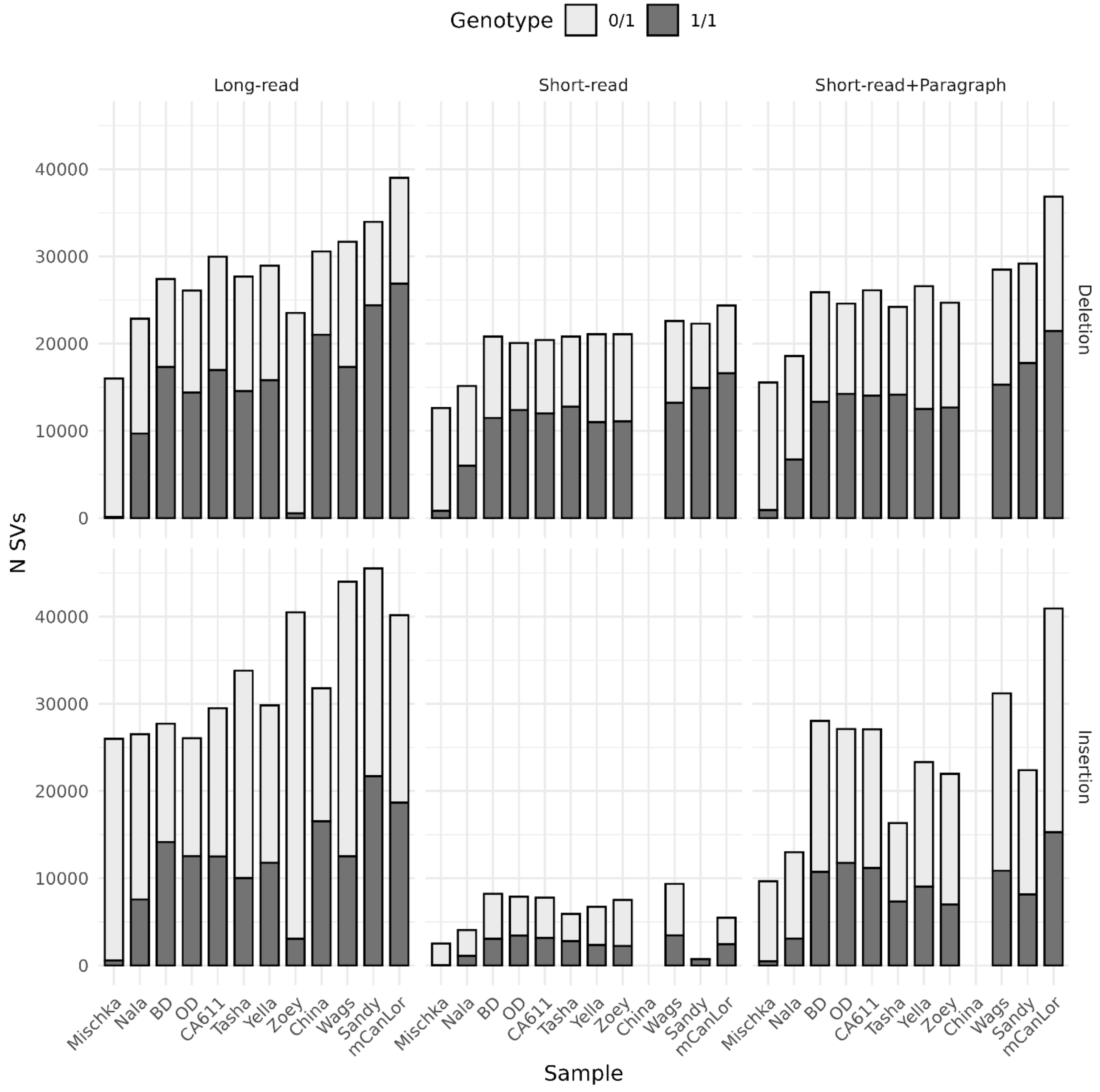


**Supplemental Figure S3 – Distribution of heterozygous and homozygous deletions and insertions by sample and datatype**

For each of the 12 samples, the total number of deletions and insertions were quantified, then bifurcated by either heterozygous (0/1, light grey) or homozygous (1/1, dark grey), and split by datatype: long-read, short-read, and short-read+Paragraph. Assembly calls were not included as they were present/absent without zygosity information.

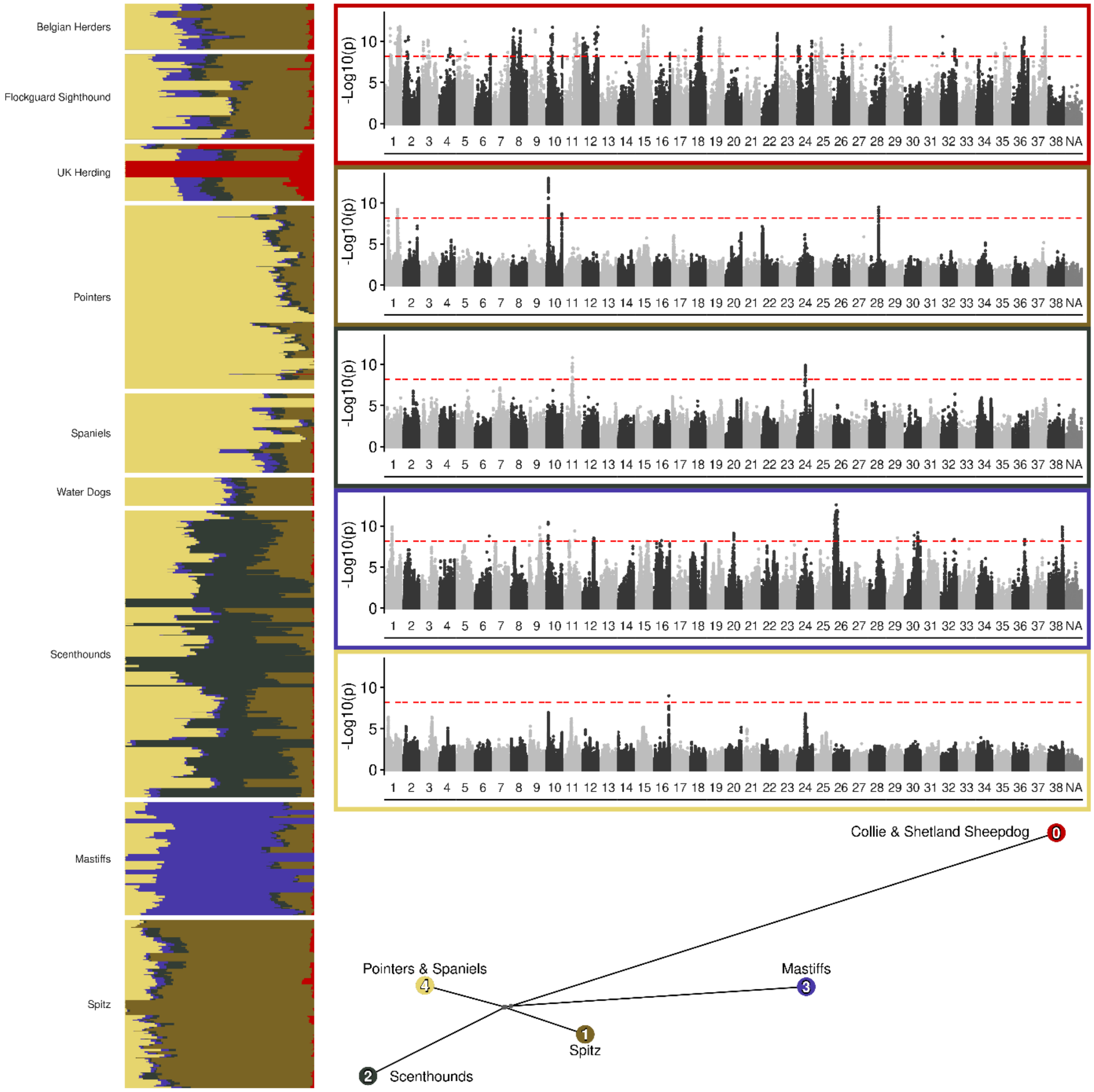


**Supplemental Figure S4 - SNP signatures of selection as detected by Ohana with five ancestral components**

Left vertical panel plots the population structure for the nine dog-breed/clade group with *K*=5 ancestral components. Manhattan plots each of the five ancestral components, exterior border color corresponds to color fill in population structure portion, horizontal dashed red-line denotes Bonferroni threshold. The bottom network tree depicts the ancestral components, labeled with the maximal dog breeds, filled color corresponding to both the Manhattan plot border and population structure.

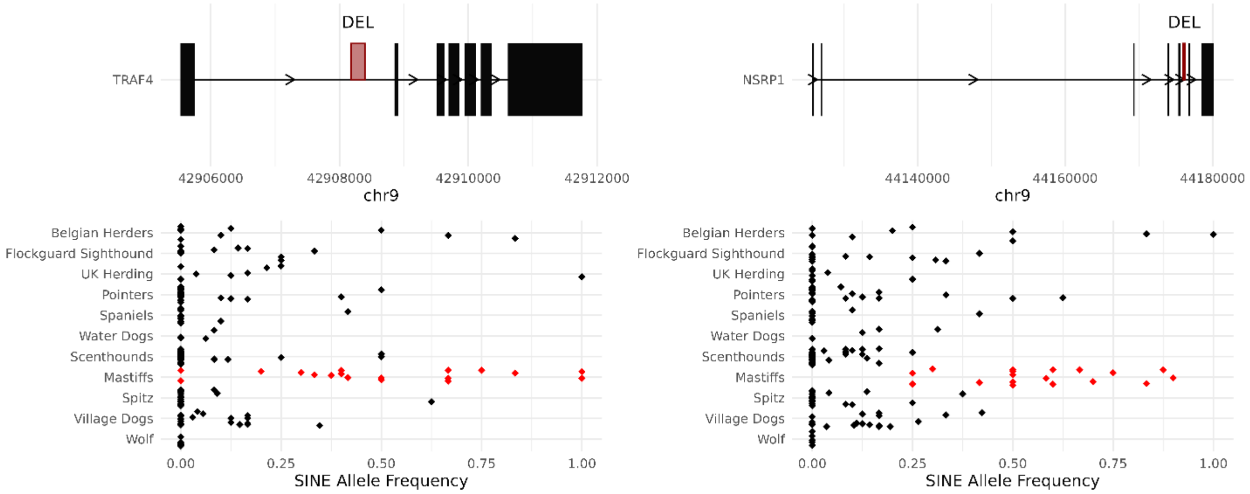


**Supplemental Figure S5 - Genomic plots SVs identified from Ohana signals of selection analysis displaying SINE frequency**

TRAF4 and NSRP1 output of Ohana, displaying the frequency of intronic SINE presence. Each panel consists of two sub panels: top displaying the specific gene within the region, highlighting the exons and region of the specific SV. The bottom panel displays the frequency of the SINE sequence presence, calculated as 1-allele frequency, by breed or geographic location. Within the bottom allele frequency panel, the red colored dots correspond to the clade selected via ohana for the associated SV.

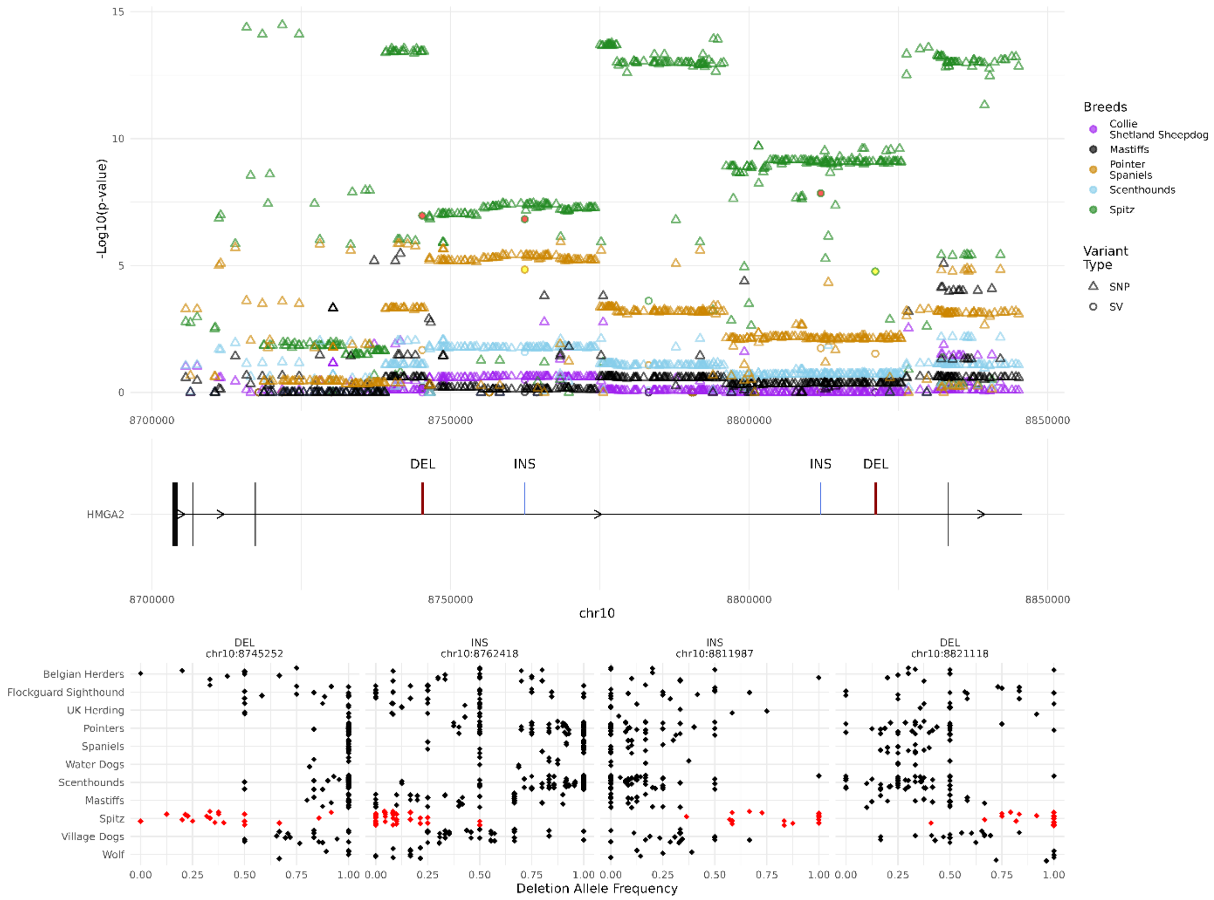


**Supplemental Figure S6 – *HMGA2* structural variants with signals of selection**

The output of Ohana, combining SNP and SV data, for the *HGMA2* gene, across the included canine breed/clade categories is depicted. Figure consists of three sub panels: the top displaying the genomic coordinates along the x-axis and the significance along the y-axis reported by Ohana, the middle displays the relevant gene model within the region, with thick boxes denoting the position of exons and the position of most significant SV given in red, the bottom displays the allele frequency by breed or geographic location for each SV. For each top panel, the color denotes the canine grouping and shape delineating variant type. Within the bottom allele frequency panel, the red colored dots correspond to the clade identified via Ohana for the associated SV selection signal. Ohana output can be found in Supplemental Table S7, and allele frequencies in Supplemental Table S8.

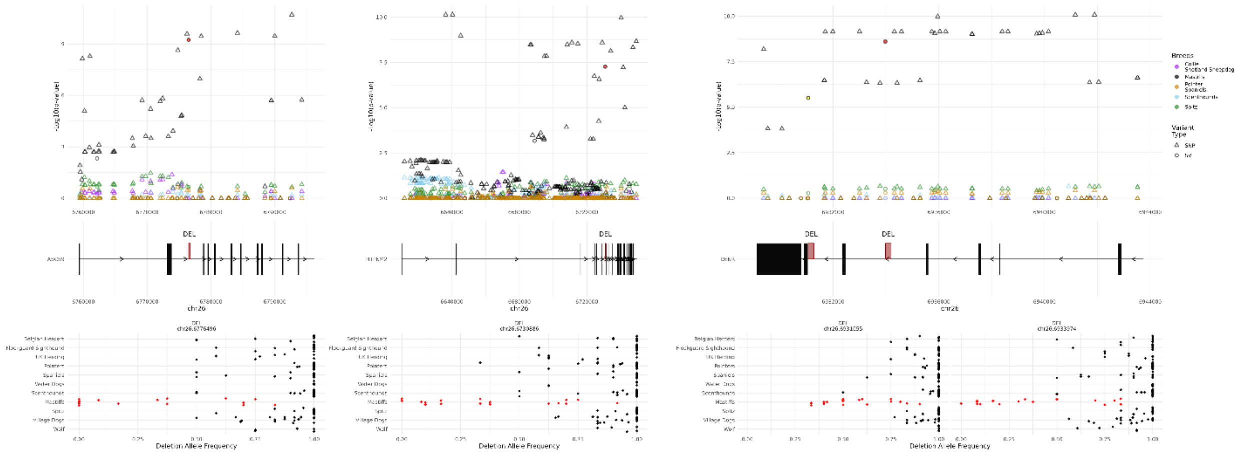


Wolf Ancestral Sequence

CTTGTGAAAGGAATAAC**AAAGAGGGG**ATGCTTGGGTGGCTCAACAGTTGA

Inserted SINE Sequence

TCTGTGCAAAGGTCCTGAGGCAGAAAGGTGCCTACTTGTGAAAGGAATAA
C**AAAGAGGGG**GGATCCCTGGGTGGCGCAGCGGTTTGGCGCCTGCCTTTGG
CCCAGGGCCGATCCCGGAGACCCGGGATCGAGTCCCACATCGGGCTCCCG
GTGCATGGAGCCTGCTTCTCCCTCTGCCTGTGTCTCTGTCTCTCTCTCTC
TCTCTGTGACTATCATGAATAAATAAATAAAATCTTTAAAAAAAAAAAAA
AAAAAAA**AAACAGGGG**ATGCTTGGGTGGCTCAACAGTTGAACATCTGCCTT

Wolf Ancestral Sequence

GATTTCCATGTG**GAAAATTGACAGCAAA**AGTGTAATGGGGAGAAGGTTGT

Inserted SINE Sequence

AAAAAAAAAAAAAAAGACTAACCCACTTTGAACTAGAACTTGATTTCCATG
TG**GAAAATTGACAGCAAA**GGGATCCCTGGGTGGCGCAGCGGTTTGGCGCC
TGCCTTTGACCCAGGGCGCGATCCTGGAGACCCAGGATCGAATCCCACGT
CGGGCTCCCGGTGCATGGAGCCTGCTTCTCCCTCTGCCTATGTCTCTGCT
TCTCTCTCTCTCTCTCTCTCTCTCTCTCTCTCTCTCTGTGACTATCATAA
ATAAATAAAAATTAAAAAAAAAGTTTAAAAAAAAAAAAAAAAAAAAA**GAAAA**
**TTGACAGCAAA**AGTGTAATGGGGAGAAGGTTGTCAGGGAGTCCT

Wolf Ancestral Sequence

AAAAATATCTTTGATATA**CAATTCCAGTTATAG**TCTTGCTGTTTGTTTTT

Inserted SINE Sequence

TTGAATTTCAAAATGGAAAAATATCTTTGATATA**CAATTCCAGTTATAG**G
GATCCCTGGGTGGCGCAGCGGTTTGGCGCCTGCCTTTGGCCCAGGGCACG
ATCCCGGAGACCCGGGATCGAATCCCACGTCAGGCTCCCGGTGCATGGAG
CCTGCTTCTCCCTCTGCCTGTGTCTCTGCCTCTCTCTCTGTCTCTGTGTG
TGACTATCATAAATAAATAAAAAAAAATTTAAAAAAAAAAAAAAAAAAAA
AAAAA**CAATTCCAGTTATAG**TCTTGCTGTTTGTTTTTTAAACAGAAATTG

Wolf Ancestral Sequence

ATGCTGTGCTTAACC**AAAAGACTTGTTAGTTTG**GCTTTTTTGGAAATACC

Inserted SINE Sequence

TGTCAGTCAGGTCCTGGCTCAGATGACTAGATAAAGCATGCTGTGCTTAA
CC**AAAAGACTTGTTAGTTTG**AGGGATCCCTGGGTGGCGCAGCGGTTTGGC
GCCTGCCTTTGGCCCAGGGCGCGATCCTGGAGACCCGGGATCGAATCCCA
CGTTGGGCTTCCGGTGCATGGAGCCTGCTTCTCCCTCTGCCTGTGTCTCT
GCCTCTCTCTCTCTCTCTCTCTCTCTCTCTCTCTCTGTGACTATCATAAATA
AATTAAAAAAAAAAAAAAAAAA**AAAAGACTTGTTAGTTTG**GCTTTTTTGGAA
ATACCAAGTATTTTTTTGG

**Supplemental Figure S7 – Ohana output and comparison of wolf sequence and inserted SINE sequence for *ABCB9, PITPNM2,* and DENR loci**

The output of Ohana, combining SNP and SV data, for three examples in their specific genomic context across the included canine breed/clade categories is depicted. Each panel consists of four sub panels: the top displaying the genomic coordinates along the x-axis and the significance along the y-axis reported by Ohana, the second displays the relevant gene model within the region, with thick boxes denoting the position of exons and the position of most significant SV given in red, the third the allele frequency by breed or geographic location for the most significant SV. For each top panel, the color denotes the canine grouping and shape delineating variant type. Within the allele frequency panel, the red colored dots correspond to the clade identified via Ohana for the associated SV selection signal. A comparison of the nucleotide sequence is given, comparing the ancestral 'empty site' represented by the wolf genome (mCanLor) and the resultant sequence with the inserted SINE sequence. For both genes/loci, the bolded text highlights the target site duplication sequence. For ease of visualization, all sequences are shown relative to the SINE forward orientation. Ohana output can be found in Supplemental Table S7, and allele frequencies in Supplemental Table S8.

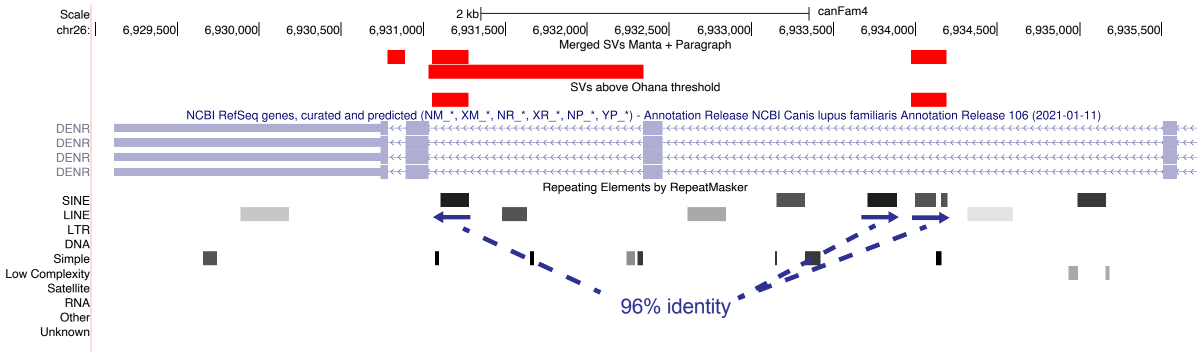


### Supplemental Figure S8 Inverted SINEC sequences in introns of *DENR*

A UCSC genome browser view of the *DENR* locus is shown. The top portion shows deletions identified and genotyped in the Dog10K collection in red. Two deletion variants were identified as evolving under selection in Mastiff clade by Ohana, as indicated. The *DENR* gene model is shown in light blue, followed by an annotation of common repeats. The two deletion variants identified by Ohana correspond to dimorphic SINEC sequence present within the introns of *DENR*. A SINEC insertion that is not variable among the samples analyzed is also located nearby. The orientations of the SINEC sequences are indicated by blue arrows. The proximal insertion, which is present in a complementary orientation, shares 96.4% (134/139 aligned positions) sequence identity with the non-variable SINEC sequence located 2,434 bp away and 95.8% (139/145 aligned positions) sequence identity with the variable SINEC sequence located 2,721 bp away.

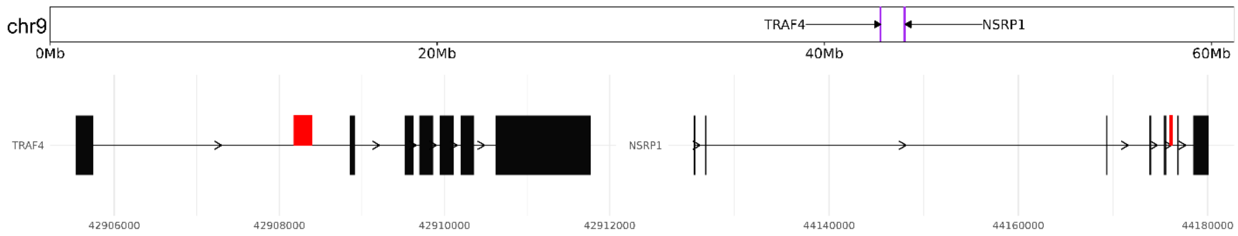


TRAF4 – Wolf Ancestral Sequence

CGGAGTGAGCCTCTCCTCTTTGCTTTAGA**AGACACACCCTTG**TGGGAGGA

TRAF4 – Inserted SINE Sequence

CGGAGTGAGCCTCTCCTCTTTGCTTTAGA**AGACACACCCTTG**GGATCCCT
GGGTGGCGCAGCGGTTTGGCGCCTGCCTTTGGCCCAGGGCGCGATCCTGG
AGACCCGGGATCGAATCCCACATCAGGCTCCCGGTGCATGGAGCCTGCTT
CTCCCTCTGCCTGTGTCTCTGCCTCCCTCTCTCTCTCTGTGTGACTATCA
TAAATAAATAAAAATTAAAAAAAAAAAAAAAAAAAAAAAAAAA**AGACACA**
**CCCTTG**TGGGAGGA

NSRP1 – Wolf Ancestral Sequence

AATAACATCACTCAAGGATGGGCATTT**AGAAAAAGAATAACA**TTAATTCA

NSRP1 – Inserted SINE Sequence

AATAACATCACTCAAGGATGGGCATTT**AGAAAAAGAATAACA**GGGGATCC
CTGGGTGGCGCAGCGGTTTGGCGCCTGCCTTTGGCCCAGGGCGCGATCCT
GGAGACCCGGGATCGAATCCCACGTCAGGCTCCCGGTGCATGGAGCCTGC
TTCTCCCTCTGCCTGTGTCTCTGCCTCTCTCTCTCTCTCTGTGACTATCA
TAAATAAATAAAAAAAAAAAAAAAAAATTTAAAAAAAAAAAAAAAAAAAA
**AGAAAAAGAATAACA**TTAATTCA

**Supplemental Figure S9 – Comparison of the wolf sequence and inserted SINE sequence for *TRAF4* and *NSRP1* loci**

Figure denotes the relative genomic position of deletions within TRAF4 and NSRP1, as identified from ohana and noted in Figure 9. Additionally, a comparison of the nucleotide sequence is given, comparing the ancestral from the wolf genome (mCanLor) and the resultant sequence with the inserted SINE sequence. For both genes/loci, the bolded text highlights the target site duplication sequence. Sequences are depicted relative to the forward orientation of each SINE.

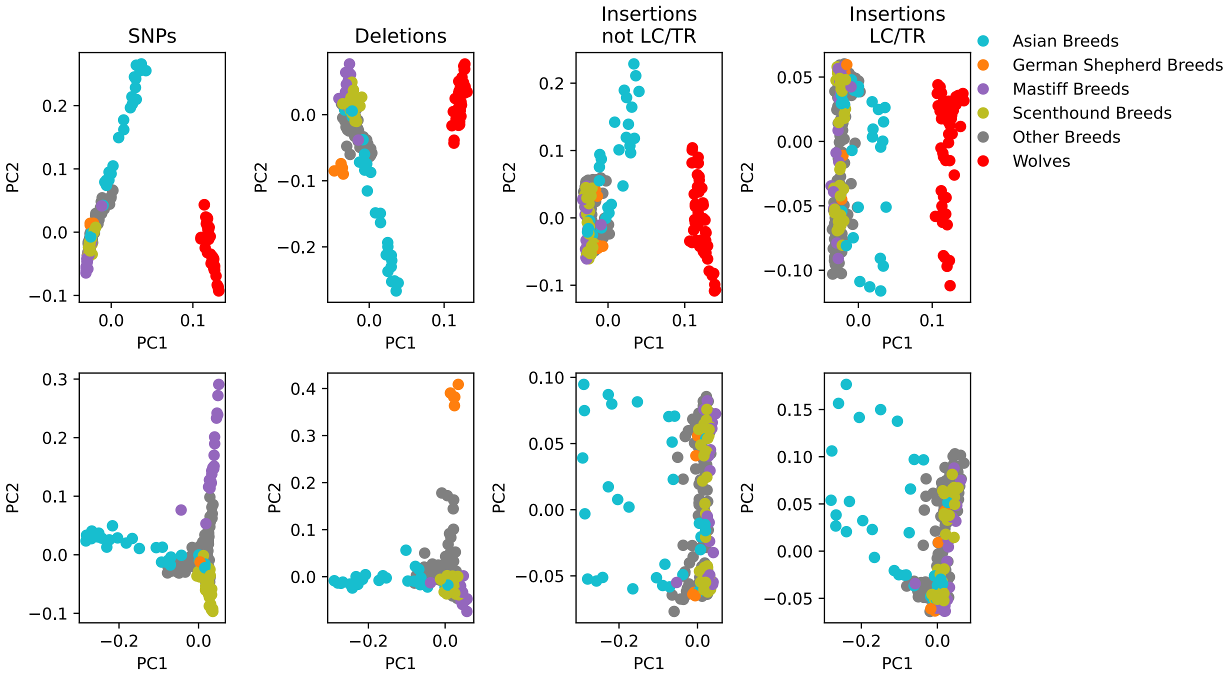


**Supplemental Figure S10 – Comparison of canine population structure measured by SNPs and SVs**

Sample relationships were assessed based on principal component analysis using different variant types. Samples were projected onto the first two components identified through an analysis of 55 wolves and one dog from each of 318 breeds (top row) or from 318 breed dogs alone (bottom row). Analysis was limited to sites that pass all genotype criteria. Loci with a missing genotype rate greater than 5%, a minor allele frequency less than 5%, or in strong linkage disequilibrium were removed. Results are shown based on SNPs (238,999 loci), deletions (21,659 loci), insertions that do not involve low-complexity sequence or tandem repeats (22,717 loci), and insertions that are low-complexity or tandem repeats (3,114 loci). In each plot wolves and breed dogs belonging to the Asian, German Shepherd, Mastiff, or Scenthound clades defined by the Dog10K Consortium are colored as indicated.
